# Endogenously generated Dutch-type Aβ nonfibrillar aggregates dysregulate presynaptic neurotransmission in the absence of detectable inflammation

**DOI:** 10.1101/2025.02.25.639746

**Authors:** Emilie L. Castranio, Merina Varghese, Elentina K. Argyrousi, Kuldeep Tripathi, Yong Huang, Akiko Asada, Linda Söderberg, Erin Bresnahan, David Lerner, Francesca Garretti, Hong Zhang, Jonathan van de Loo, Cheryl D. Stimpson, Ronan Talty, Charles Glabe, Efrat Levy, Minghui Wang, Marjan Ilkov, Toshiharu Suzuki, Kanae Ando, Bin Zhang, Lars Lannfelt, Brigitte Guérin, William D. Lubell, Shai Rahimipour, Dara L. Dickstein, Sam Gandy, Ottavio Arancio, Michelle E. Ehrlich

**Author notes:** Correspondence or, Michelle E. Ehrlich, M.D., Professor of Neurology, Pediatrics, Genetics, and Genomic Medicine, Sam Gandy, M.D., Ph.D., Professor of Neurology and Psychiatry, Icahn School of Medicine at Mount Sinai, Once Gustave L. Levy Place, Box 1138, New York NY 10029. Contributed equally to supervision (Sam Gandy, Ottavio Arancio, Michelle E. Ehrlich).

## Abstract

**Background:** *APP^E693Q^* (“Dutch”) transgenic mice develop aging-related learning deficits and accumulate endogenously generated nonfibrillar aggregates of Aβ (NFA-Aβ) and APP α-carboxy terminal fragments. NFA-Aβ correlates with synaptic loss and memory deficits more closely than does fibrillar Aß.

**Methods:** We assessed the physiological, transcriptomic, ultrastructural, histological, and metabolic changes associated with the accumulation of NFA of Dutch Aβ in brains of *APP^E693Q^* mice.

**Results:** Aging-related accumulation of NFA-Aβ in *APP^E693Q^* mice was revealed by A11 immunohistochemistry and cyclic D,L-α-peptide-FITC microscopy. Presynaptic termini of *APP^E693Q^* mice developed physiological abnormalities in post-tetanic potentiation, synaptic fatigue, synaptic vesicle replenishment, and an aging-related reduction in mitochondrial complex I activity. Single-cell RNA sequencing showed that excitatory neurons exhibited an altered transcriptomic profile involving “protein translation” and “oxidative phosphorylation”.

**Discussion:** Accumulation of NFA-Aß alters neuronal metabolism but does not activate inflammation. Depletion of all forms of Aβ may be required to eliminate Aβ toxicity with anti-amyloid antibodies.

## Introduction

Synaptic dysfunction is believed to be a key pathological hallmark of Alzheimer’s disease (AD) that begins gradually and parallels clinical cognitive decline (*1–4*). Many studies have demonstrated that the burden of fibrillar amyloid-β (Aβ) proteinopathy does not correlate with synaptic loss (*5–7*) or with cognitive status (*8*). Instead, some investigators have proposed that brain levels of synaptotoxic nonfibrillar aggregates of Aβ (NFA-Aβ) including Aβ oligomers and protofibrils correlate more closely with synaptic loss (*9, 10*). Synaptic NFA-Aβ has been detected in AD brains, and NFA-Aβ levels in human brain synaptosomes can differentiate subjects with early AD from cognitively intact controls with AD pathology (*11–13*). *In vitro*, synthetic NFA-Aβ peptides can cause inhibition of long-term potentiation (LTP) in hippocampal slices (*14, 15*), while *in vivo* injections of NFA-Aβ into the cerebral ventricles of healthy rats leads to cognitive deficits (*16*). Studies using a transgenic mouse model of AD have demonstrated that neuronal loss in the hippocampus correlates with NFA-Aβ levels but not with amyloid plaque burden (*17*). In 2010, we reported a transgenic mouse model overexpressing Dutch mutant human amyloid precursor protein (*APP^E693Q^*) on the pan-neuronal Thy1 promoter (*18–25*). This mutation within the Aβ domain substitutes a glutamine for glutamic acid residue, thereby disrupting fibril assembly and favoring the formation of NFA-Aβ. The *APP*^E693Q^ mutation causes hereditary cerebral hemorrhage with amyloidosis - Dutch type (HCHWAD) (*19*). The neuropathology of HCHWAD shows prominent cerebrovascular amyloidosis and hemorrhage as well as diffuse parenchymal Aβ deposition (*20*). Although the *APP^E693Q^* (or “Dutch”) transgenic mouse failed to recapitulate the full neuropathology of either AD or HCHWAD, we were struck by the aging-related development of learning behavior impairment that was observed in the absence of detectable fibrillar Aβ. Analysis of the Dutch mouse brain revealed that this learning behavior impairment was directly related to the brain levels of NFA-Aβ (*18, 21*), an observation supported by imaging data in the current paper using epitomic histochemistry with anti-prefibrillar oligomer-specific antibody A11 (*26*) and NFA-Aβ-detecting cyclic peptide-FITC fluorescence microscopy (*22*). The Dutch mice also exhibit detectable cerebrovascular Aβ deposition and traces of perivascular iron (presumably due to the extravasation of red blood cells) without overt hemorrhage.

Differential exon usage was also prominent in the Dutch mouse entorhinal cortex and the PSAPP (*APP^KM670/671NL^/PSEN1^Δexon9^*) mouse dentate gyrus. The transcriptomes from the entorhinal cortex and the dentate gyrus of Dutch mice were also enriched in markers of glutamatergic neurons, synaptic transmission, and protein translation. To study transcriptomics in the Dutch mouse hippocampus, single-cell RNA sequencing (scRNA-seq) was employed and revealed dysregulation of pathways for protein translation, oxidative phosphorylation, and the electron transport chain. Direct measurements of electron transport chain function revealed changes in activity of mitochondrial complex I but not in that of complexes II, III, or IV.

In summary, a multi-disciplinary analysis of the Dutch mouse has now revealed alterations in synaptic physiology, gene expression, and protein translation in excitatory neurons, as well as changes in the transcriptomic profile and mitochondrial morphology and activity. Moreover, the high sensitivity of scRNA-seq has enabled neuroinflammation to be ruled out as a major contributor to pathogenesis in the hippocampi of Dutch mice. The present studies are among few focusing on endogenous generation of NFA-Aβ without the concurrent presence of potential confounds from either Aβ fibrils or synthetic peptides. These findings support the notion that fibrillar Aβ is not necessary for the aging-related learning deficit observed in Dutch mice. These data may have important implications for the genesis of cognitive impairment in humans. The clinical response to anti-amyloid antibody therapy may be dampened due to the persistence of toxic NFA-Aβ conformers that go undetected in contemporary analysis of fluid biomarkers and in amyloid fibril positron emission tomography (PET) imaging. Depletion or neutralization of NFA-Aβ may be required for complete elimination of clinically relevant Aβ toxicity.

## Materials and Methods

### Experimental animals

Male and female 2- to 24-month-old mice were used in this study. All animals were group housed in microisolator cages under a 24-hour light/dark cycle and given *ad libitum* access to food and water. The Dutch mice used in the study express the human *APP* gene harboring the Dutch mutation (*APP*^E693Q^) under control of the *Thy1* promoter (*18*). All animal procedures were conducted in accordance with the National Institute of Health Guidelines for the Care and Use of Experimental Animals and were approved by the Institutional Animal Care and Use Committees at the Icahn School of Medicine at Mount Sinai LA10-00462.

### Dot blot assay

The cerebral cortices of mouse brain samples were thawed and lysed by adding 2x volume of lysis buffer [5.0 mM NaCl, 5.0 mM EDTA, 1.0 mM dithiothreitol (DTT), and protease inhibitor mixture in 25 mM Tris buffer, pH 7.5] containing 0.5 mm Zirconium Oxide Beads (ADV-ZrOBO5, ORNAT, Israel), homogenized with a tissue grinder motor pestle (Alex-Red, Israel) on ice, and the mixtures were centrifuged at 4°C at 14,000 rpm for 15 minutes. The total protein in each sample was quantified by Nano-Drop absorbance quantification at 280 nm.

For dot-blot analysis, equal amounts of samples of diluted proteins (100 ng/μL) were directly applied in duplicate onto nitrocellulose membrane and dried at room temperature for 30 minutes. The membrane was then blocked with 5% bovine serum albumin (BSA) in tris-buffered saline (TBS) (pH 7.4) containing 0.1% (v/v) Tween 20 (TBST). Blots were incubated overnight at 4°C with: 6E10 mouse monoclonal antibody (1:1000); rabbit polyclonal antibody recognizing prefibrillar oligomers, A11 (1:5000); rabbit monoclonal antibody OC recognizing fibrillar oligomers (1:1000); anti-murine-α-tubulin antibody (1:5000) in TBST containing 5% BSA (all antibodies used are listed in **Supplementary Table 1**). The membranes were washed with TBST three times and incubated for 1 hour at room temperature with horseradish peroxidase (HRP) conjugated either to an anti-mouse IgG (1:10000) or anti-rabbit IgG (1:5000) in 5% BSA in TBST. The blots were washed with TBST and developed using the enhanced chemiluminescence (ECL) reagent kit (Pierce) in Amersham Imager 680 machine. Images were then used to analyze the mean of integrated densities of each blot using ImageJ software (National Institutes of Health, USA) (*27*).

### Immunohistochemistry (IHC) and CP-2-FITC incubation

All staining was performed in paraformaldehyde (PFA)-fixed 30 μm thick sections of Dutch and wildtype mouse brains. For A11 immunohistochemistry and cyclic D,L-α-peptide-2 (CP-2)-FITC fluorescence, free-floating sections were washed with 0.1 M phosphate buffer saline (PBS) in 0.1% Triton (PBST) three times for 15 minutes and blocked with 2% bovine serum albumin (BSA) for 1 hour. Sections were incubated with a rabbit polyclonal antibody A11, recognizing Aβ prefibrillar oligomers (1:2500) in antibody buffer (PBST containing 2% BSA) overnight at 4°C. Sections were washed and incubated with Alexa Fluor 680 secondary goat anti-rabbit antibody (1:1000 Abcam) in antibody buffer for 1 hour at room temperature. Sections were then incubated with fluorescein-5-isothiocyanate (FITC)-conjugated CP-2 [0.15 mg/mL in 2% dimethyl sulfoxide (DMSO)] for 3 hours at room temperature. Sections were washed for 10 min in 1% PBST and kept for 15 min to dry. Sections were then mounted on super frost glass slides, embedded with Fluro-Gel II mounting medium containing DAPI (4’,6-diamidino-2-phenylindole dihydrochloride) (Electron Microscopy Sciences, Hatfield PA), cover-slipped, and stored in the dark for microscopic imaging. Immunofluorescence images containing the hippocampus, cortex, and thalamus were obtained using the 20x objective of a Leica DM6000 microscope (Leica Microsystems, Wetzlar, Germany), coupled to a controller module and a high sensitivity 3CCD video camera system (MBF Biosciences, VT) and an Intel Xeon workstation (Intel). All images were analyzed by the Image J program (*27*), and one section from each mouse (from a total of 5 mice in each group) were taken for analysis. The average intensity was calculated from the area of the hippocampus, cortex, and thalamus.

### Brain tissue homogenization and Aβ ELISAs

Fresh frozen brain tissue (cerebrum) was homogenized at a 1:5 tissue:buffer ratio, using a Precellys Evolution homogenizer (Bertin Technologies, Montigny-Le-Bretonneux, France) to sequentially extract TBS, TBS with 0.5% Triton X-100 (TBS-T) and formic acid (FA) soluble fractions of Aβ. After homogenization in TBS, samples were centrifuged at 16,000× *g* for 1 h at 4 °C (TBS_16K_). For a subset of experiments, a fraction of this supernatant was further centrifuged at 100,000× *g* followed by collection of the supernatant (TBS_100K_). Homogenization of the remaining tissue pellet was repeated according to the same procedure with TBS-T, then followed by FA. Brain extracts were analyzed with Aβ1-40 and Aβ1-42 ELISAs, as previously described (*28*). In brief, 96-well half-area plates were coated over night with 50 ng/well of anti-Aβ40 (custom production, Agrisera) or anti-Aβ42 (700254, Thermo Fisher Scientific, USA), then blocked with 1% BSA in PBS for 3 h at RT. TBS_16K_, TBS-T and FA brain extracts were diluted in ELISA incubation buffer (PBS, 0.1% BSA, 0.05% Tween) and incubated over night at 4°C, followed by detection with 0.5 µg/ml biotinylated 3D6 and HRP-conjugated streptavidin (Mabtech AB, Nacka, Sweden). Signals were developed with K Blue Aqueous TMB substrate (Neogen Corp., Lexington, KA, US). Plates were developed and read with a spectrophotometer at 450 nm. The TBS and TBS-T extracts were also analyzed for soluble Aβ aggregates using two different sandwich ELISAs. The first, based on 3D6 as both capture and detection antibody, allows for detection of Aβ aggregates from the size of a dimer, but not for monomeric Aβ. The second preferentially detects larger soluble Aβ aggregates as it utilizes the Aβ protofibril selective antibody mAb158 for capture and 3D6 for detection (*29*). Ninety-six-well half-area plates were coated over night with 3D6 (50 ng/well) and blocked with 1% BSA in PBS for 3 h at RT. Brain extracts were diluted in ELISA incubation buffer (PBS, 0.1% BSA, 0.05% Tween) and incubated over night at 4 °C, followed by 3D6 biotin detection and development as above.

### Slice preparation, electrophysiological recordings, and analysis

For electrophysiological experiments, hippocampi from 7- to 13-month-old Dutch and wildtype mice were excised on ice and coronal hippocampal slices (400 μm) were cut with a tissue chopper. Slices were transferred to a recording chamber where they were allowed to recover for 2 hours at 29°C and perfused with artificial cerebrospinal fluid (ACSF) containing (124.0 mM), KCl (4.4 mM), Na_2_HPO_4_ (1.0 mM), NaHCO_3_ (25.0 mM), CaCl_2_ (2.0 mM), MgCl_2_ (2.0 mM), and glucose (10.0 mM). ACSF was bubbled with carbogen (95% O_2_ and 5% CO_2_) and perfused at a flow rate of 2 ml/minute.

Field excitatory post-synaptic potentials (fEPSP) were measured after stimulating the Schaffer collateral fibers with a bipolar tungsten electrode placed at the CA3 and recording at the *stratum radiatum* of CA1 with a glass pipette filled with ACSF. Basal synaptic transmission was evaluated by plotting the relationship between increased voltages (5 V to 35 V) and evoked fEPSP responses, generating an input-output (I-O) curve. Recordings were measured at approximately 35% of the maximum evoked response as determined by the I-O curve. For LTP recordings, a 30-minute baseline was recorded prior to theta-burst stimulation (4 pulses at 100 Hz, with the bursts repeated at 5 Hz and 3 tetani of 10-burst trains administered at 15-second intervals), responses were recorded for 2 hours after tetanization and measured as fEPSP slope expressed as percentage of baseline. Paired-pulse facilitation (PPF) was induced by pairing two pulses at intervals of 15, 20, 30, 40, 50, 100, 200, 300, 500 and 1000 ms. PPF was expressed as the ratio between the second and the first fEPSP slopes. Post-tetanic potentiation (PTP) was induced by the above-described theta-burst stimulation protocol in presence of the NMDA receptor blocker D-APV (50 µM). Synaptic fatigue (SF) and replenishment were measured by 100 Hz stimulation followed by 7 stimuli spaced at 125 msec from each other in the presence of D-APV (50 µM).

Results were analyzed in pClamp 11 (Molecular Devices) and fEPSP slopes were compared by 2-way ANOVA for repeated measures for the I-O curve, LTP, PPF and PTP recordings. For statistical comparison of synaptic fatigue and replenishment, the rates of exponential growth/decay (k) extra sum-of-squares F test was used to test whether k differed between data sets vs. the data sets sharing one k-value. Statistics were performed using GraphPad Prism 9 software. Data were expressed as mean ± SEM. The level of significance was set at p<0.05.

### Perfusions and tissue processing for electron microscopy and array tomography

For electron microscopy (EM) and array tomography synapse studies, 12-month-old Dutch and wildtype mice were anesthetized intraperitoneally, and transcardially perfused with 1% PFA in phosphate-buffered saline (PBS; pH 7.4) followed by 4% PFA with 0.125% glutaraldehyde in PBS as described previously (*24, 30, 31*). The brains were carefully removed, hemisected, postfixed overnight at 4°C in 4% PFA in PBS with 0.125% glutaraldehyde for synapse and immunogold analyses, or by 2% PFA with 2% glutaraldehyde in PBS for presynaptic vesicle analysis, and sectioned on a Vibratome (Leica VT1000S, Bannockburn, IL) into 250-μm thick sections. All sections were stored at 4°C in PBS with 0.1% sodium azide until use. Coronal sections encompassing the CA1 region of the hippocampus were prepared for EM as reported (*24, 31*). For synapse and immunogold analyses, slices were cryoprotected in graded phosphate buffer/glycerol washes at 4°C, and manually microdissected to obtain blocks containing the CA1 region. The blocks were rapidly freeze-plunged into liquid propane cooled by liquid nitrogen (-190°C) in a Universal cryofixation System KF80 (Reichert-Jung) and subsequently immersed in 1.5% uranyl acetate dissolved in anhydrous methanol at -90°C for 24 hours in a cryosubstitution unit (Leica). Block temperatures were raised from -90°C to -45°C in steps of 4°C/hour, washed with anhydrous methanol, and infiltrated with Lowicryl resin (Electron Microscopy Sciences) at -45°C. The resin was polymerized by exposure to ultraviolet light (360 nm) for 48 hours at -45°C followed by 24 hours at 0°C. For presynaptic vesicle studies, tissue blocks were washed overnight in 0.1 M sodium cacodylate buffer, placed in 1% tannic acid/0.1 M cacodylate buffer solution at room temperature for one hour and washed 3 times for 10 minutes each in 0.1 M sodium cacodylate. The samples were then incubated for 1 hour at room temperature in 2% osmium tetroxide/double distilled water (ddH_2_O), washed again in 0.1 M sodium cacodylate 3 times for 10 minutes then subjected to a graduated ethanol dehydration series. After dehydration, the tissue samples were infiltrated with Spurr Low Viscosity Embedding Kit (Sigma Aldrich) and polymerized for 12 hours at 70°C. For both EM preparations, block faces were trimmed, ultrathin sections (90 nm) were cut with a diamond knife (Diatome) on an ultramicrotome (Reichert-Jung), and sections were collected on formvar/carbon-coated nickel slot grids (Electron Microscopy Sciences).

### Post-embedding immunogold labeling for EM

Immuno-gold labeling was performed using a previously optimized method (*32*). Briefly, the EM sections were treated with 0.1% sodium borohydride in 50 mM glycine prepared in Tris buffer for 6 minutes in the dark, to remove aldehydes. Sections were then subjected to six quick rinses and two 10-minute washes in TBS. The sections were incubated for 30 min in blocking buffer consisting of 2% human serum albumin (HSA) in TBS with 0.1% Triton X-100. After three short washes in TBS, the sections were incubated in anti-glutamate receptor 2 & 3 rabbit polyclonal antibody (Millipore AB1506, diluted 1:100 in 2% HSA in TBS) at room temperature overnight with gentle shaking. Following six quick rinses and two 5-minute washes in TBS, the grids were incubated in blocking buffer for 20 minutes, washed three times, and incubated for 90 minutes at room temperature in goat anti-rabbit IgG conjugated to 10-nm gold particles (Aurion 25362), diluted 1:40 in 2% HSA and 5 mg/ml polyethylene glycol in TBS. The grids were rinsed six times and washed for 5 minutes in TBS and then in deionized water. Imaging was done using an H-7000 electron microscope (Hitachi High Technologies America, Inc.) with an AMT Advantage CCD camera (Advanced Microscopy Techniques) at 15,000x. The electron micrographs were adjusted for brightness and sharpness with Photoshop CS5 (Adobe Systems).

### Quantifying synapse density, ultrastructure, and distribution of immunogold particles at mGluR-2/3-labeled synapses

All EM analyses were conducted by an experimenter blinded to the genotypes of the mice. For synapse quantification and ultrastructural analysis, nine sets of serial images across the same set of 5 consecutive ultrathin sections were taken from the *stratum radiatum* of CA1 and imported into Adobe Photoshop (version CS5). To obtain a stereologically unbiased population of synapses for quantitative morphologic analysis, we used a disector approach on ultrathin sections as described (*24, 30, 31, 33*). Briefly, all axospinous synapses were identified within the first and last 2 images of each 5-section serial set and counted if they were contained in the reference image (layers 1 and 4) but not in the corresponding look-up image (layers 2 and 5). To increase sampling efficiency, the reference image and look-up image were then reversed; thus, each animal included in the current study contributed synapse density data from a total of 18 disector pairs. The total volume examined was 11.317 μm^3^, and the height of the disector was 180 nm. Axospinous synapse density (per μm^3^) was calculated as the total number of counted synapses from both images divided by the total volume of the disector. The criteria for inclusion as an axospinous synapse included the presence of synaptic vesicles in the presynaptic terminal and a distinct asymmetric PSD separated by a clear synaptic cleft. For a synapse to be scored as perforated it had to display two or more separate PSDs.

To measure structural parameters such as spine head diameter and PSD length, we identified all axospinous synapses in the middle portion of three serial sections. The synapse and spine head were followed through layers 2-5. In each layer, the PSD length (if present) and the spine head diameter at its widest point parallel to the synapse were measured. PSD area was determined by summing the PSD lengths through all sections and multiplying by section thickness (90 μm). This same process was repeated for unique synapses in layer 4 (reference layer 5), following the synapses from layer 4 to layer 1. Each of these synapses were also classified as non-perforated or perforated. A total of 734 total synapses (341 from Dutch mice and 393 from wildtype) were analyzed in this manner. An average of 81 synapses per animal and a minimum of 71 synapses per animal were sampled. For each unique synapse in layer 2 in each series (as identified for PSD length and spine head analysis above), the synapse was followed through layers 2 though 5 and the number of immunogold particles in each of the following areas were counted: active zone (presynaptic, within 45 nm of the membrane across from the PSD; region A), synaptic cleft (region B), PSD (region C), peri-synaptic (peri—PSD, within 200 nm of the PSD on the post-synaptic spine head; region D), postsynaptic membrane (>200 nm away from the PSD; region E), and postsynaptic cytoplasm (region F). A cutoff of 45 nm away from the membrane was used to differentiate between receptors at the membrane and receptors in the cytosol.

For presynaptic vesicle analysis, 100 synapses per animal were analyzed from at least 30 different images taken at a magnification of 8,000x. PSD length, maximum spine head diameter, and cytosolic area of the presynaptic terminal were measured using Adobe Photoshop 2023. Vesicles in the presynaptic terminal were labeled with a dot in the center of the vesicle, and a line was drawn over the PSD. An algorithm was written in Python that measured the shortest distance between each vesicle (represented by the dot in the center) and its corresponding PSD (represented by the manually drawn line over the PSD) and reported a list of measurements for all vesicles within an adjustable threshold distance. For each synapse, the total number of vesicles was determined with output from this algorithm, along with their individual distances from the PSD. The vesicles were divided into bins of 10 and 50 nm for comparison of spatial distribution within the presynaptic terminal. Additionally, vesicles within 10 nm of the PSD were classified as docked, based on a 17 nm average minimum diameter of vesicles in our samples (*34–36*).

### scRNA-seq and data analysis

For the scRNA-seq experiments, fresh hippocampi were dissected from 12- to 13-month-old Dutch and wildtype mice, and the tissue was dissociated into a single-cell suspension following the 10X Genomics sample preparation protocol. We pooled the hippocampi from 3 mice of the same gender, genotype, and age to reach the target concentration of 2000 cells/µl. The single cell cDNA libraries were generated using a V3 Chromium kit (10X Genomics) and sequenced using a NovaSeq S4 (Illumina).

The raw sequencing data were processed using the standard Cell Ranger pipeline to generate unique molecular identifier (UMI) count matrices. Starting from UMI matrices, we first performed quality control (QC) by removing low-quality cells with either too few (< 200) or too many (> 4000) genes detected, retaining 61,642 cells after filtering. We then removed insufficiently detected genes by keeping 20,548 genes expressed in more than one cell. After QC, we obtained 5,266 ∼ 10,949 cells per individual, and an average of 618 unique genes per cell per individual. We performed a clustering analysis using a well-established scRNA-seq data integration workflow based on R package Seurat (v3) (*37*). Briefly, the UMIs data were first normalized by sequencing depth and log-transformed using the LogNormalize method implemented in Seurat. The 2,000 most variable gene features were identified, scaled, and centered. The scRNA-seq data across all sequencing libraries were combined based on the canonical correlation analysis (CCA) integration method available from Seurat. Next, dimensional reduction was performed using principal component analysis (PCA) based on the 2,000 most variable genes. The top 25 principal components (PCs) collectively explaining more than 90% of the variance were selected for calculating 2D dimensional reductions by Uniform Manifold Approximation and Projection for Dimension Reduction (UMAP) (*38*). The same top 25 PCs were also used to compute the nearest neighbor graph and the subsequent cell clusters with the Louvain algorithm implemented in Seurat (*37*). This clustering analysis resulted in 14 clusters (named as Clusters 0 – 13, sorted by cluster size in a decreasing order) at a clustering resolution of 0.1. These clusters contained 267 to 20,702 cells each.

### Cluster cell type annotation

For each cluster, we first interrogated the expression patterns of known gene markers to annotate clusters into major cell-types: inhibitory neurons (*Gad1*, *Gad2*), excitatory neurons (*Slc17a7*), astrocytes (*Aqp4*), oligodendrocytes (*Mog*), microglia (*Csf1r*, *Tyrobp,* and *Trem2*), oligodendrocyte progenitor cells (*Vcan* and *Pdgfra*), endothelial cells (*Flt1*), and pericytes (*Vtn* and *Pdgfrb*). Next, we calculated *de novo* cluster signatures by comparing the cells in a cluster against the cells of the other clusters using Wilcoxon rank sum test in Seurat. We defined cluster marker genes as those upregulated by at least 1.2-fold and with Bonferroni adjusted p-value less than 0.05. To assist the annotation of cell type of each cluster, we overlapped the *de novo* cluster marker gene signatures with a large-scale collection of cell type markers curated from over 1,054 single-cell experiments (*39*), with p-value significance of the overlaps computed by a hypergeometric test.

### Cluster-specific differential gene expression and functional enrichment analysis

DEGs between Dutch and wild type sample groups in each cluster were identified using MAST (*37, 40*) implemented in Seurat. Similarly, DEGs between males and females, and DEGs between Dutch and wild type in each sex group were identified using MAST. DEGs were determined at the cutoff of Bonferroni-corrected p-value ≤ 0.05. Functional enrichment of DEGs with MSigDB gene annotation collections was examined by a hypergeometric test. Results with Benjamini-Hochberg (BH) correction p-value ≤ 0.05 were considered statistically significant.

### Microglia subclustering analysis

We studied a subset of microglial cells from the full data matrix. PCA was performed and then the top PCs collectively explaining more than 90% of the variance were selected for calculating dimensional reduction by UMAP (*38*) as well as for clustering analysis using Seurat (*37*). To test if the microglia subclusters represent known subtypes, we computed single cell signature scores (*41*) corresponding to homeostatic, disease-associated microglia (DAM), type I IFN-induced (IFN) and MHC-expressing (MHC) microglia based on signature genes curated by Chen and Colonna (*42*).

### Quantifying presynaptic mitochondrial numbers and morphology

For analysis of mitochondria, 12 series of 9 consecutive 90-nm thick sections were selected from each mouse using a systematic-random approach for EM imaging. All mitochondrial analyses were performed by the same experimenter, blinded to the group designation of subjects as described (*43*). For each series of 9 consecutive sections, the middle fifth section was used as a reference, and all pre-synaptic axonal boutons containing a mitochondrion were identified. Each of these boutons was followed throughout the series, and the number, length and morphology of mitochondria were recorded. Mitochondria were classified as either straight or curved. Mitochondria that were bent at least 90° were identified as curved, while all other mitochondria were classified as straight.

### Statistics

Data sets from the dot blot assays were analyzed using a one-way ANOVA followed by Tukey’s multiple comparisons test. Data sets from the A11 immunohistochemistry and CP-2 cyclic peptide-FITC (CP-2-FITC) (*22*) fluorescence microscopy were analyzed using a two-way ANOVA followed by Sidak’s multiple comparisons test. Data sets were analyzed using a two-tailed Student’s t-test. Threshold for significance was set at α = 0.05. Distribution of mGluR-2/3 immunogold particles was analyzed using repeated measures ANOVA. Data are represented as mean ± SD. Statistical analyses were carried out using Prism software (version 9.2.0, GraphPad Software, San Diego, CA).

### Data Availability

The scRNA-seq data (cell ranger processed count matrices) are in a Synapse repository (accession syn52752404, with direct link https://www.synapse.org/#!Synapse:syn52752404/files/).

## Results

### Dutch mice show increased levels of nonfibrillar aggregates with aging

Aging-dependent investigation of the Dutch mice revealed that 12 months is the first age at which deficits attributable to NFA-Aβ can be detected (*18*). To determine the involvement of NFA-Aβ in brain changes, multiple analyses were conducted at different ages, before and after the appearance of NFA-Aβ. Epitomic characterization was performed on cortical tissue from 2- and 24-month-old Dutch mice. Interestingly, this characterization revealed a significant increase with aging in the A11/6E10 ratio (ratio of prefibrillar oligomer content to total Aβ content; F (3,16) =17.05; p<0.0001), but no such aging-related change in the OC/6E10 ratio (ratio of fibrillar oligomer content to total Aβ content; F (3,16) =2.557; p=0.0916) in Dutch mice (**Figure 1, Supplementary Figure 1**). Furthermore, brains were analyzed for the presence of protofibrils (PF) using the homogenization protocol described in Tucker *et al.* (*44*) and assayed for PF levels in the TBS soluble fraction using a mAb158-mAb158 sandwich MSD assay employing an in-house synthetic protofibril as calibrator. Both assays showed very low levels of Aβ PF. In the mAb158 sandwich assay, Aβ PF levels were around 100 pM, levels indistinguishable from those in homogenates of wildtype mice (**Supplementary Figure 1**). No PF were detected in the cerebellum. Levels of PF in Dutch mice were approximately 10- to 100-fold lower than those measured in Tg-APP^ArcSwe^ and Tg2576 (depending on age analyzed). In addition, Aβ_38/40/42_ fragments were analyzed in the TBS insoluble pellet fraction after dissolution in formic acid and neutralization. Levels of Aβ PF were again very low, consistent with the absence of parenchymal plaques in Dutch mice.

**Figure 1.**
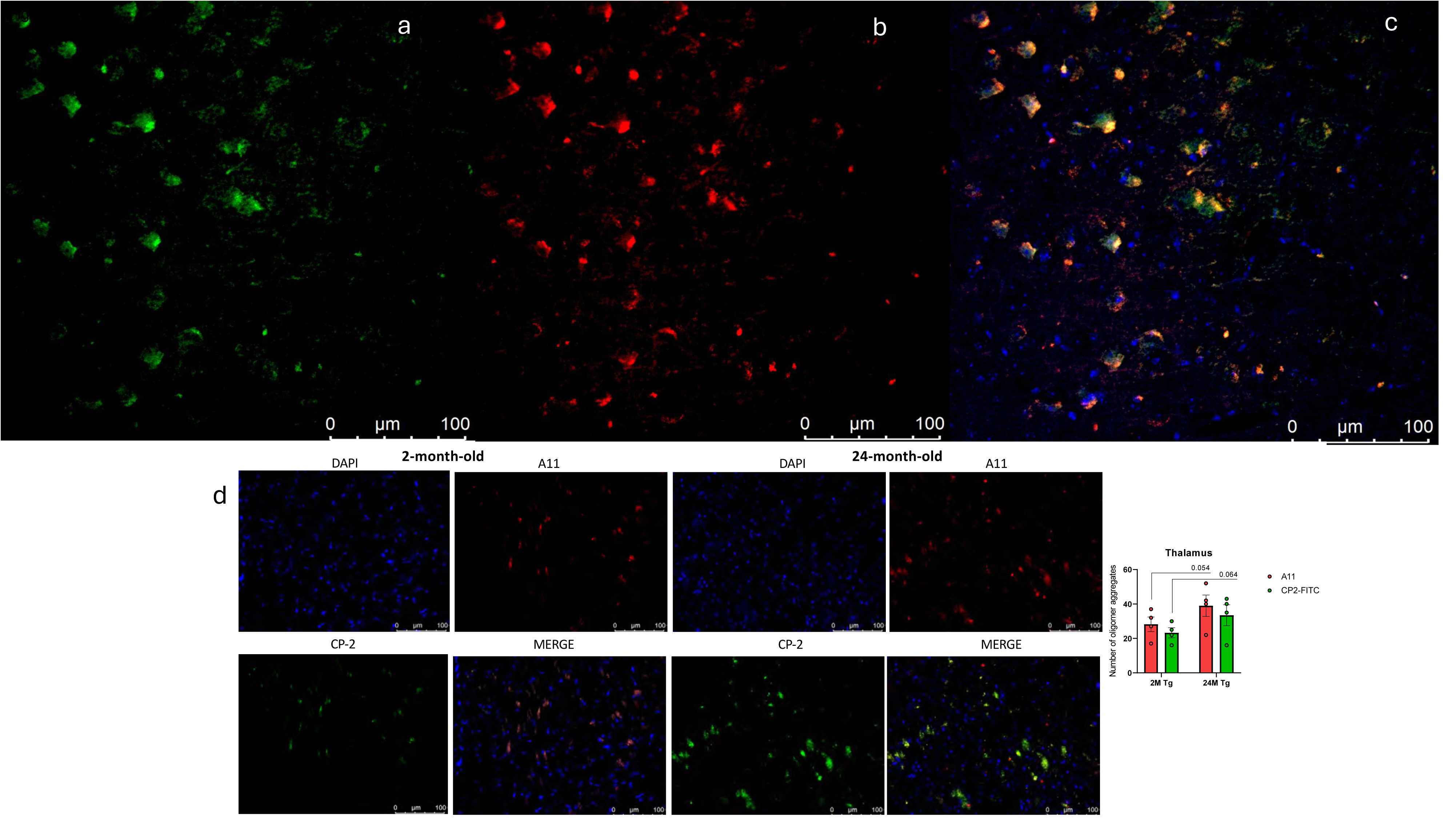
Aging-related accumulation of NFA-Aβ in the cerebral cortex (a)-(c) and thalamus (d) of APP^E693Q^ transgenic mice revealed by double labeling with A11 epitomic immunohistochemistry and CP-2 cyclic peptide-FITC fluorescence microscopy. Representative images from 2-month-old and 24-month-old Dutch mice showing fluorescence microscopy using DAPI, A11 staining with AlexaFluor-680 secondary antibody (red), and CP-2 cyclic peptideFITC (CP-2-FITC, green)(*22*). Upper right panel is a merge of A11 and CP-2-FITC images. Scale bar = 100 µm. Quantification of A11-immunoreactive and CP-2-FITC-immunoreactive NFA-Aβ aggregates in 2- and 24-mo Dutch shown at bottom right. ** p < 0.01.

The elevation of NFA-Aβ levels in the cortex, thalamus, and hippocampus of Dutch mice was visually and morphologically confirmed using a combination of immunohistochemistry using A11 antibody and fluorescence microscopy employing cyclic D,L-α-peptide-2 (CP-2)-FITC as a fluorescein-tagged NFA-Aβ-detecting ligand (*22*) (**Figure 1, Supplementary Figures 1-3**). The hypothesis that CP-2 and A11 bind to similar or identical domains on NFA-Aβ was confirmed by cross-competition blocking experiments in which pre-incubation of brain slices with CP-2 peptide prevented the subsequent binding of the prefibrillar oligomer-specific antibody A11 (**Supplementary Figure 4**). Furthermore, the interaction of CP-2 with NFA-Aβ has been characterized (*45–49*): CP-2 was demonstrated (1) to stabilize Aβ dimers and trimers using a photoinduced cross-linking of unmodified proteins (PICUP) assay (*45–48*) and (2) to bind 10 nm particles typical of NFA-Aβ (*49*). In the current study, CP-2 exhibited similar K_d_ values for both fibrillar and nonfibrillar forms of Aβ in surface plasmon resonance (SPR) assays (**Supplementary Methods**): CP-2 interacted with Aβ fibrils and NFA-Aβ with K_d_ values of 6 and 4 mM, respectively.

As shown in **Supplementary Table 2**, there were no important male-female differences in the levels of NFA Aβ in either Dutch or Dutch PS1 mice. Among the neurophysiological data, the differences in replenishment were statistically significant regardless of whether they were analyzed by sex or whether data from different sexes were pooled for analysis. For fatigue and PTP analyses, only one sex was significant, but this was probably because of the disparity of sample sizes rather than a sex-specific effect. There were insufficient numbers of individual sexes studied by EM to calculate sex-specific mitochondrial size reliably.

### Dutch mice showed no significant impairment in basal synaptic transmission and long-term synaptic plasticity in the hippocampus

The relationship between synaptic input strength and the resulting electrical response, the so-called “input-output relation” and long-term potentiation (LTP) were tested in 7- to 7.5-month-old and 10.5- to 11-month-old Dutch mice. The two genotypes displayed similar input-output relationships and LTP in both age groups, suggesting that basal synaptic transmission at the CA1 region of the hippocampus and long-term synaptic plasticity at the Schaffer collateral pathway were unimpaired in Dutch mice **(Figure 2A, 2B, 2C, 2D**).

**Figure 2.**
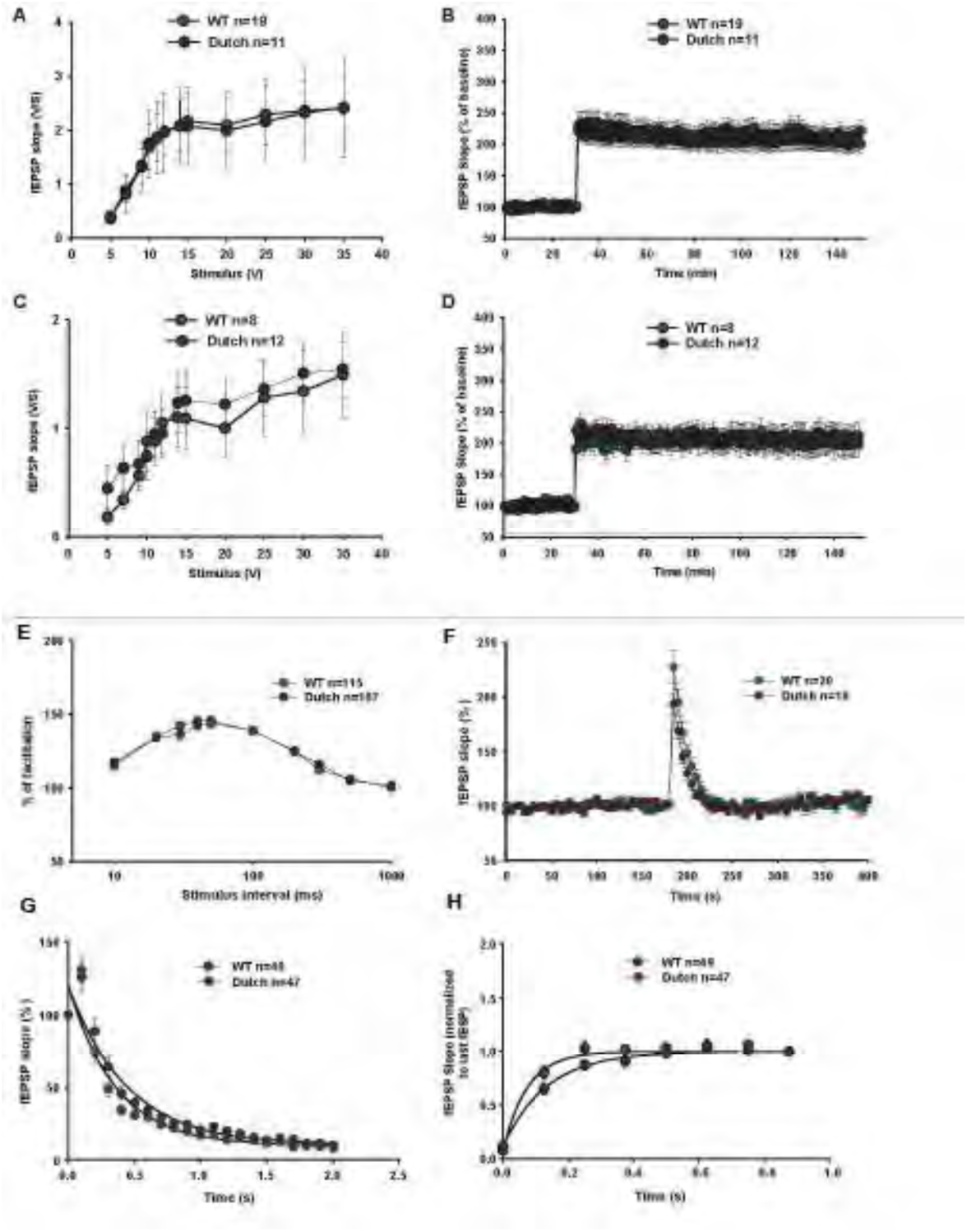
Dutch mice do not exhibit impairment in basal synaptic transmission or LTP but showed impairment of certain measurements of short-term synaptic plasticity. **(A, B)** Basal neurotransmission **(A)** and LTP **(B)** are unimpaired in hippocampal slices from Dutch mice as compared to slices from wildtype littermates at the age of 7 to 7.5 months (F _(1,28)_ =0.0026, p=0.959 and F _(1,28)_ =0.219, p=0.644, respectively). **(C-D)** Slices from Dutch mice do not exhibit impairment of basal neurotransmission **(C)** and LTP **(D)** at the age of 10.5 to 11 months compared to wildtype mice (F _(1,18)_ =0.116, p=0.737 and F _(1,18)_ =0.017, =0.899, respectively). **(E)** Dutch mice showed no impairment in paired-pulse facilitation at the age of 11 to 13 months old (F _(1, 220)_ =0.06364, p=0.8011; 2-way ANOVA). **(F)** Post-tetanic potentiation was significantly reduced in slices from 11-to 13-month-old Dutch mice compared to wildtype (F _(1,36)_ =4.771, p=0.036; 2-way ANOVA for the first 6 points after potentiation). **(G)** Significant differences were observed in synaptic fatigue when comparing slices from Dutch mice and wildtype littermates (k=2.247±0.1622 for wildtype and k=2.782±0.1634 for Dutch, p=0.015, k: rate of exponential decay). **(H)** The rate of synaptic vesicle replenishment following a 100 Hz stimulation was found to be significantly increased in hippocampal slices from Dutch mice as compared to slices from wildtype littermates (k=7.509±0.7277 for wildtype, k=13.64±1.682 for Dutch, p<0.0001, k: rate of exponential growth).

### Dutch mice exhibited defects in PTP, SF, and replenishment of the vesicle pool after depletion

We assessed several mechanisms related to function of the presynaptic terminal. Both 11- to 13-month-old Dutch mice and their wildtype littermates showed the same response in paired-pulse facilitation (PPF) **(Figure 2E)**, a type of short-term plasticity that is most likely dependent upon the build-up of Ca^2+^ in the presynaptic neuron due to the close pairing of two action potentials. Accumulation of Ca^2+^ in the presynaptic neuron results in increased neurotransmitter release and subsequently increased response after the arrival of the second action potential (*50*). Accordingly, facilitation is no longer present when a long-time interval (1 second) is introduced between the two stimuli. Both wildtype and Dutch mice showed increased facilitation at short time intervals, and no difference was observed between the two genotypes. In contrast to PPF, Dutch mice showed decreased post-tetanic potentiation (PTP) compared to wildtype mice **(Figure 2F)**. PTP is performed in the presence of D-2-amino-5-phosphonovalerate (D-APV) that blocks NMDA receptors, thereby preventing the onset of NMDA-dependent postsynaptic mechanisms that might interfere with measurement of plasticity. Due to the DAPV, the fEPSP slope declines back to baseline a few seconds after stimulation. PTP is another form of short-term plasticity and reflects the enhanced release of neurotransmitter due to the increased levels of Ca^2+^ at the presynaptic terminal during the tetanic stimulation (*51*).

We also studied the effect of the Dutch mutation on the efficacy of presynaptic mechanisms by examining synaptic fatigue (SF) and replenishment, after depletion, of the vesicle pool. Both mechanisms appeared to differ between wildtype and Dutch mice. Specifically, the SF showed a faster rate of decay (**Figure 2G**) and the replenishment phase had a higher growth rate in the Dutch compared to wildtype mice (**Figure 2H**). The decline in the slope that is observed during SF is thought to represent the depletion of the readily releasable pool (RRP) of neurotransmitter that resides in vesicles at the presynaptic cells. Replenishment after depletion with high frequency stimulation depicts the refilling process of the RRP. Mechanistically, both SF and replenishment have been suggested to depend on accumulation of residual Ca^2+^ at the presynaptic terminal (*50, 51*).

### Dutch mice had larger PSD area on small mGluR-2/3^+^ synapses in the CA1 region

Changes in mGluR-2/3^+^ excitatory neurons were examined at the ultrastructural level in 12-month-old mice. Comparable percentages of mGluR-2/3^+^ synapses among total, non-perforated, and perforated synapses in the CA1 were observed in Dutch and wildtype mice using immunogold EM against mGluR-2/3 (**Figure 3A-C**). Moreover, Dutch and wildtype mice exhibited no difference in the total and perforated synapses (**Figure 3D, F**). Higher postsynaptic density (PSD) area of non-perforated mGluR-2/3^+^ synapses was detected in the Dutch compared to wildtype mice (**Figure 3E**), both of which had comparable distributions of mGluR-2/3^+^ particles per PSD area (**Figure 3G-I**).

**Figure 3.**
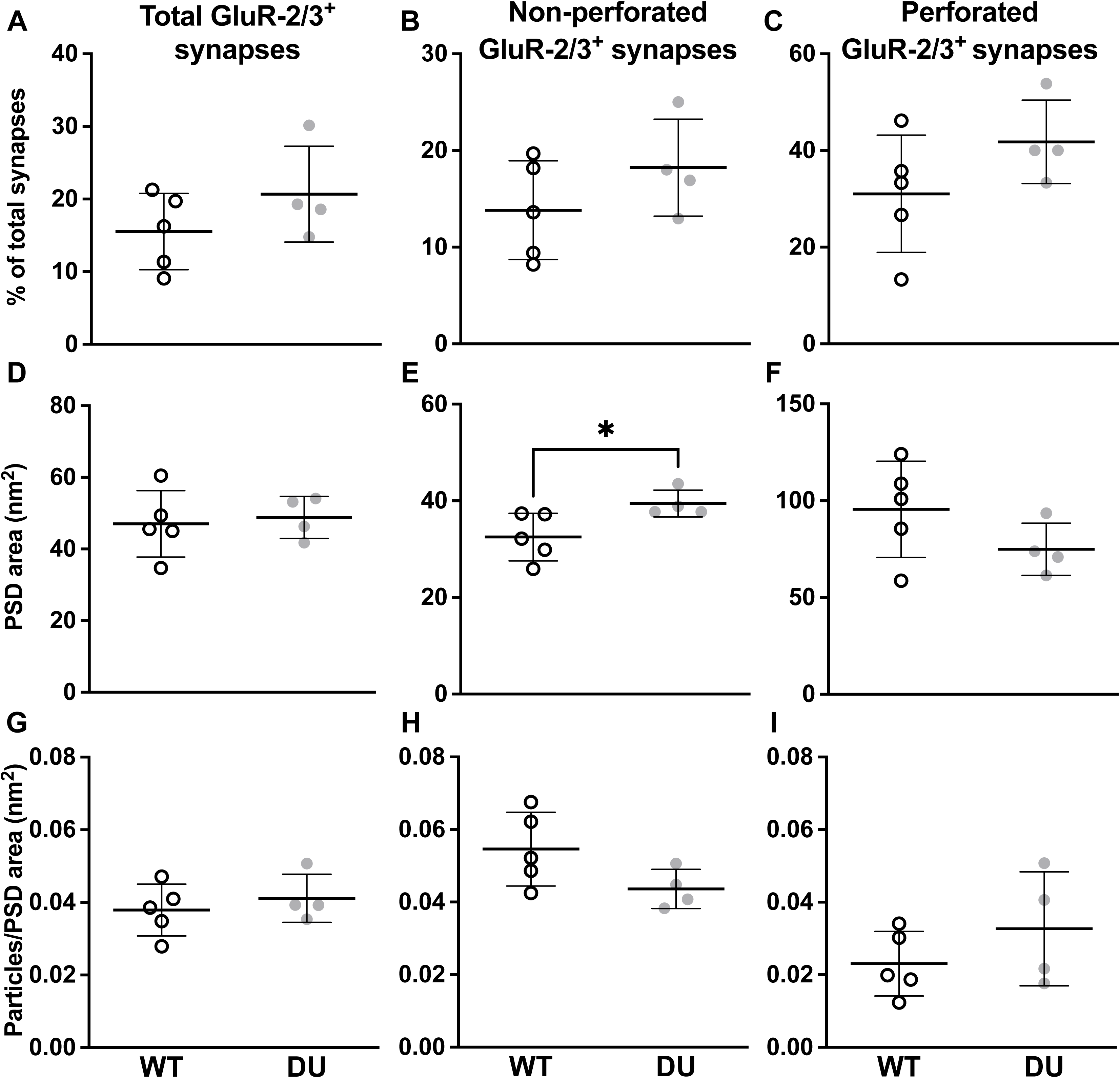
Distribution and ultrastructure of mGluR-2/3-labeled synapses were comparable in the CA1 of 12-month-old Dutch and wildtype mice. mGluR-2/3-labeled synapses were defined as having ≥1 immunogold particle located in the PSD or synaptic cleft. **(A-C)** No significant differences were observed between Dutch and wildtype mice in percentage of mGluR-2/3-labeled total, perforated, and non-perforated synapses, respectively. **(D-F)** Average PSD area of mGluR-2/3labeled synapses for all synapses, non-perforated synapses, and perforated synapses, respectively. Dutch mice had larger PSD area of mGluR-2/3-labeled non-perforated synapses than wildtype (p= 0.04, by unpaired t test), whereas perforated synapses or total synapses were comparable between the two groups. **(G-I)** No significant difference was observed in density of immuno-gold particles in mGluR-2/3-labeled synapses, calculated as number of immuno-gold particles/ PSD area, for all synapses, non-perforated synapses, and perforated synapses, respectively. Data are mean ± SD from n = 5 wildtype, and n = 4 Dutch mice.

### Dutch mice showed no change of hippocampal mGluR-2/3 expression nor of mGluR-2/3+ synapse density

Comparable levels of mGluR-2/3 protein were observed in Western blots of the insoluble membrane fraction from the hippocampus of 12-month-old Dutch and wildtype mice (**Supplemental Figure 5A**). Considering that Western blots may not detect regional differences, array tomography and immunofluorescence were used to analyze mGluR-2/3 puncta after staining serial sections of CA1 neurons with Neurotrace (**Supplemental Figure 5B**). Comparable densities in the CA1 of Dutch and wildtype mice were observed in 3D reconstructions of mGluR-2/3 synaptic puncta by array tomography (**Supplemental Figure 5C**, F _(4,3)_ = 1.476, p = 0.7802). Quantification of the total mGluR-2/3 puncta output yielded an average tissue density (puncta/μm^3^) of 0.322 ± 0.112 for Dutch and 0.275 ± 0.087 for wildtype mice. Both genotypes exhibited comparable puncta size ranges, and no significant difference in average mGluR-2/3 puncta size (**Supplemental Figure 5D**; F _(1,3)_ = 0.751, p = 0.538) per percentile per sample. Similarly, neither the density nor size of GluN1 puncta were altered in Dutch mice (**Supplemental Figure 5E-F**; F _(4,3)_ = 1.431, p = 0.8003 for density and F _(1,3)_ = 0.116, p = 0.950 for puncta size).

### Dutch mice had no change in synaptic distribution of mGluR-2/3+ in the CA1

Comparable distributions of mGluR-2/3 were observed by immunogold labeling and electron microscopy (EM) at the synapse (**Supplemental Figures 6A-C**), pre- and postsynaptic compartments (**Supplemental Figure 6D**), presynaptic active zone, synaptic cleft, peri-synaptic area, postsynaptic membrane, and postsynaptic cytoplasm **(Supplemental Figure 6E**) in 12-month-old Dutch and wildtype mice.

### Dutch mice exhibited no change in vesicle size nor number of docked vesicles but show a trend towards fewer vesicles in the reserve pool at presynaptic terminals in the CA1

The size, number, and distribution of vesicles were quantified within the presynaptic terminal of 12-month-old mice. No differences were observed in synaptic vesicle sizes at the presynaptic terminal nor the number of docked vesicles at the synapses of the active zone in Dutch and wildtype mice (**Supplemental Figure 7A-C, E, G**). A decline in the number of synaptic vesicles was found in the reserve pool beyond 300 nm from the active zone of Dutch compared to wildtype mice but did not reach statistical significance and appeared to be driven by vesicles in large synapses (**Supplemental Figures 7D and 7H**). The hippocampi of Dutch and wildtype mice exhibited no differences in the levels of the postsynaptic density protein of 95kDa (PSD95) and of the presynaptic marker synaptophysin (**Supplemental Figure 7I**).

### Dutch mice display shared gene expression profiles across multiple cell types

Employing brains from Dutch mice alongside sex- and age-matched wildtype littermates, scRNA-seq was performed. Hippocampi were collected from mice at 12 months, an age characterized by established cognitive deficits and high levels of NFA-Aβ and α-CTF in Dutch mice (*18*). After quality control filtering (**Supplemental Figure 8**), 61,642 (32,661 Dutch and 28,981 wildtype) cells were isolated from thirteen distinct clusters which were identified based on the expression of well-known cell-type-specific markers (**Figures 4, 5A-B**, **Supplemental Tables 3-4**). Pseudo-bulk differential expression analysis was used to identify gene expression changes within each cell type. For further analysis (C0-6), clusters of interest were identified based on differentially expressed genes (DEGs; adj. p-value cutoff <0.05) (**Figure 5C-D**, **Supplemental Figures 9-12, Supplemental Table 5**).

**Figure 4.**
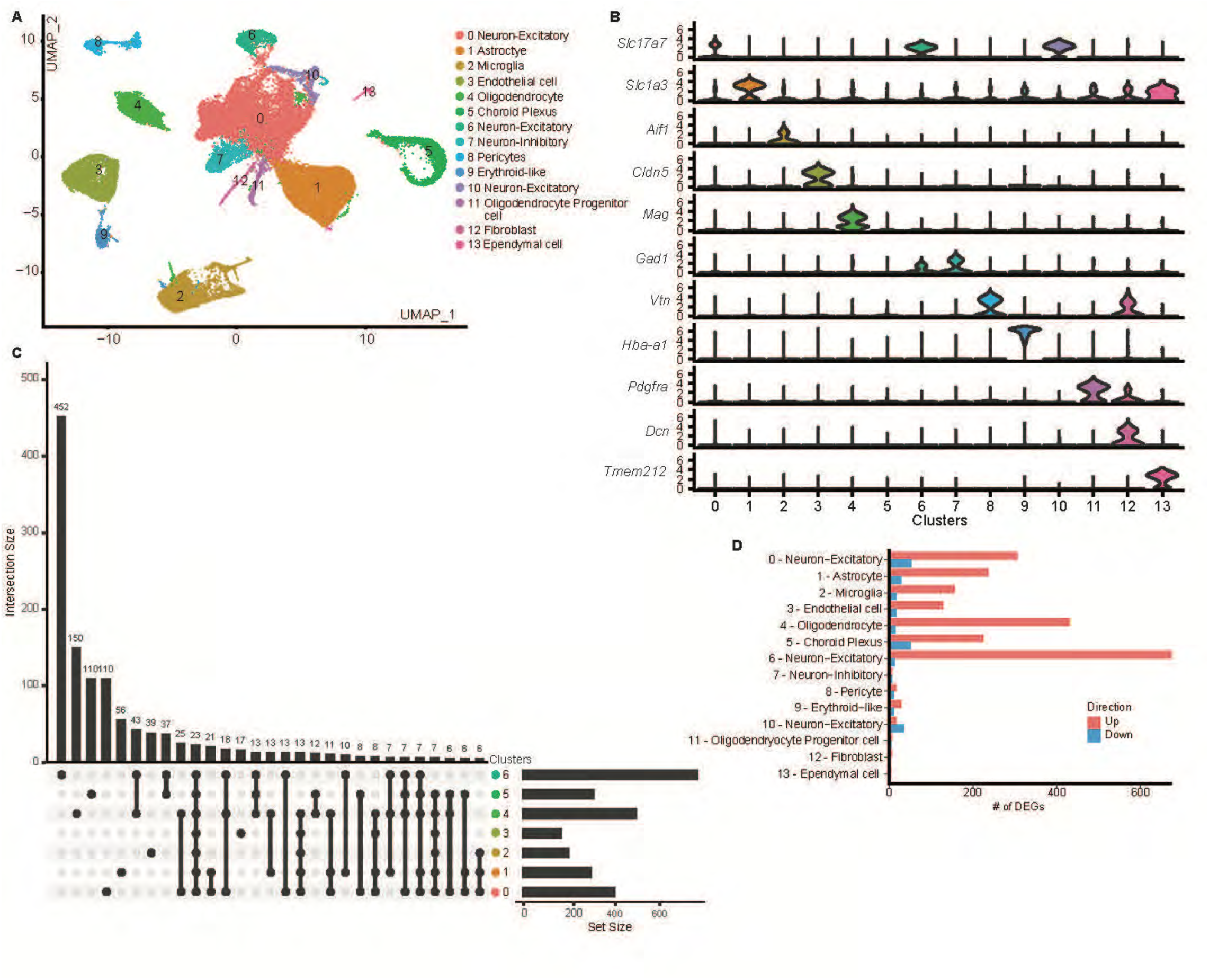
Transcriptional responses in Dutch mouse brains. **(A)** scRNA-seq of hippocampal tissues from 12-month-old male and female Dutch and wildtype mice (n=2 samples per sex per genotype with 3 mice pooled per sample). UMAP plot showing 14 distinguishable clusters (0-13). **(B)** Violin plots illustrating expression of marker genes for each cell type. **(C)** UpSet plot highlighting the overlapping and unique genes between the clusters. **(D)** Bar plot showing number of DEGs per cluster by directionality.

**Figure 5.**
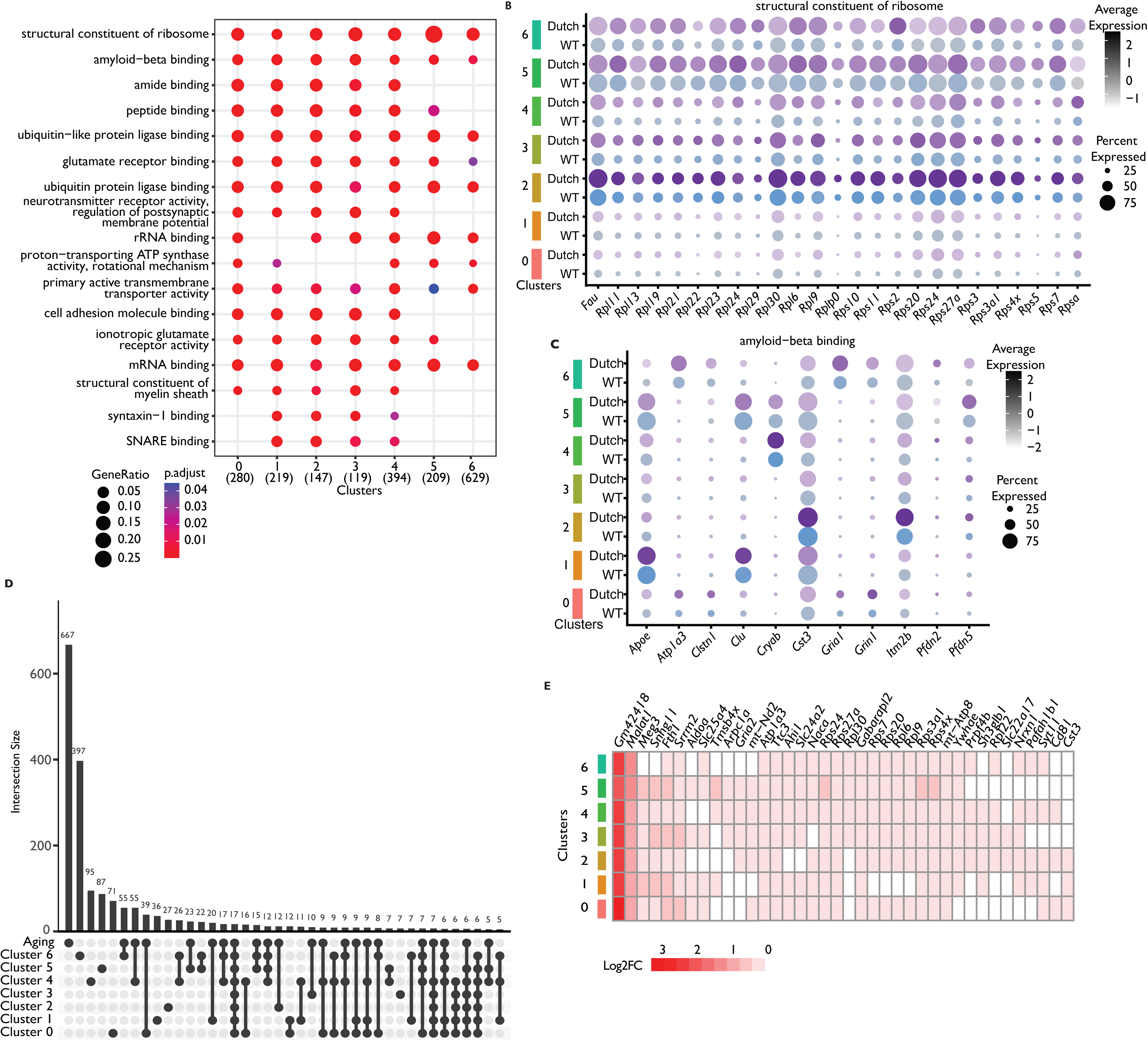
Overlapping transcriptomic profiles highlight processes impacted in multiple cell types. **(A)** Dot plot highlighting shared GO term enrichment across clusters. **(B-C)** Dot plot of genes associated with shared GO terms **(B) “**structural constituent of ribosome” and **(C) “**amyloid-beta binding”, split into Dutch and wildtype mice across all clusters. Wildtype mice are in blue, Dutch mice are in purple. **(D)** UpSet plot comparing overlap of cluster DEGs (adj. p-value <0.05) with previously defined aging gene list. **(E)** Heatmap of log2FC of genes shared with aging gene list.

Faced with considerable overlap in DEGs, the clusterProfiler (*52*) approach was employed to compare biological themes and identify the Gene Ontology (GO) term enrichment shared among the gene clusters. Among the shared terms were “structural constituent of ribosome” (example genes include: *Rpl23, Rpl30, Rpl37a, Rpl6, Rpl9, Rps24*) (**Figure 6A-B**), and “amyloid-beta binding” (*Apoe, Atp1a3, Clu, Cst3)* (**Figure 6A, C**). Across clusters, the shared DEGs and GO terms were similar to a previously identified “aging signature” in 21- to 22-month-old mouse brain (*53*), in which ribosomal protein genes, long noncoding RNA genes (e.g., *Malat1*) as well as ribosome-related pathways and processes, all were significantly affected across multiple cell types compared to that of 2-to 3-month-old counterparts. Across clusters, 453 DEGs were found to overlap with the aging-related gene list (**Figure 6D**) from which 435 genes displayed the same directionality across all clusters (389 upregulated and 46 downregulated, **Figure 6E**) as reported (*53*).

**Figure 6.**
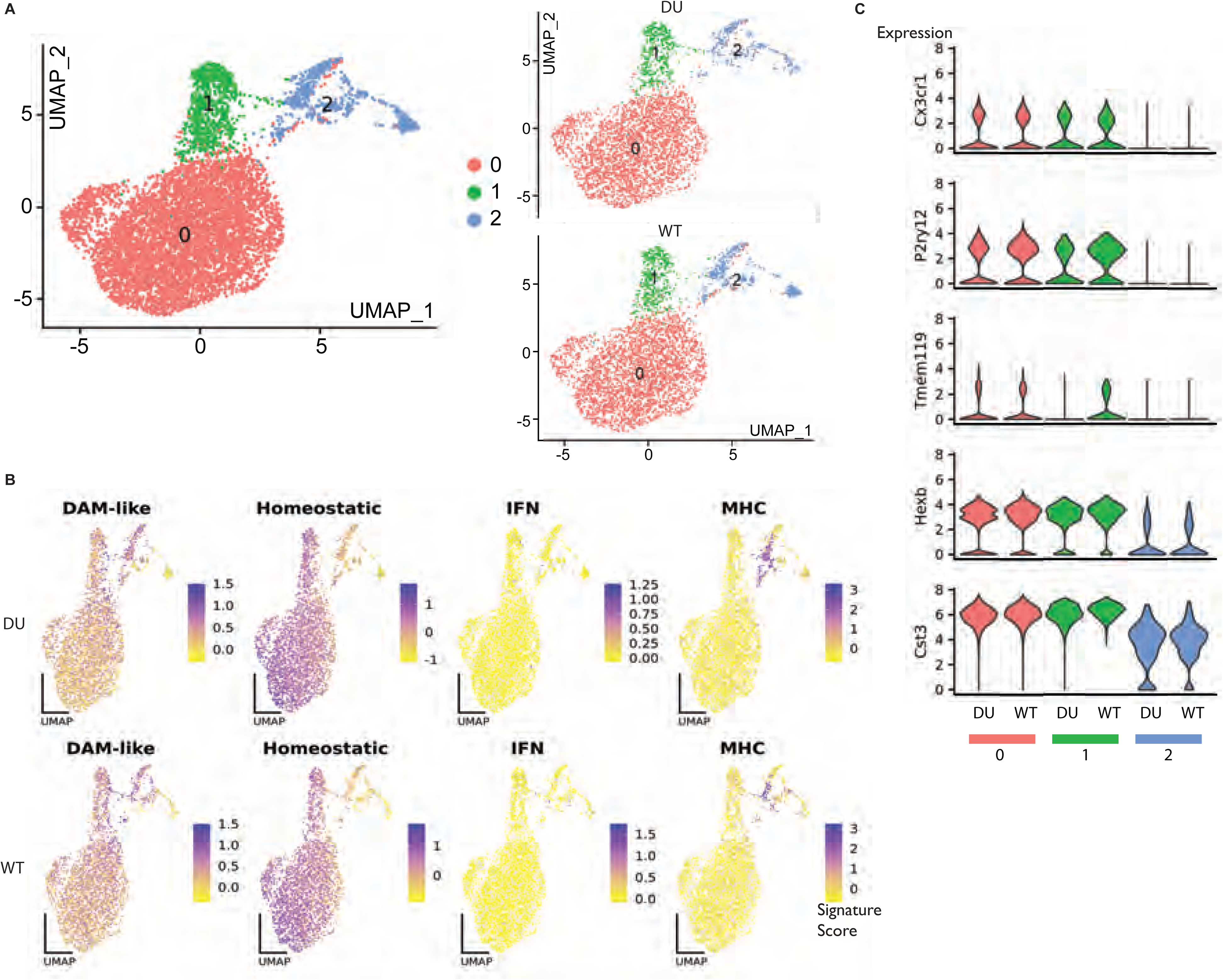
Microglia transcriptomic profiles are similar between Dutch and wildtype mice. **(A)** UMAP plot showing sub-clustering analysis results for Cluster 2 microglia, identifying 3 subpopulations of microglia with UMAP plot split by genotype to the right. **(B)** UMAP plots showing expression of previously identified microglial signatures; DAM(*33*), homeostatic, IFN and MHC(*34*). **(C)** Violin plots showing expression across subclusters by genotypes of selected homeostatic microglial genes.

### Dutch mice do not show obvious differences in microglial gene expression

In many Aβ-depositing mouse models of AD, microglia show prominent transcriptional responses and increased microgliosis with conspicuous peri-plaque localization (*54, 55*). Furthermore, the microglial immune response to NFA-Aβ has been implicated as an early, key step in AD pathogenesis. To investigate the microglial responses in more detail, we performed subclustering analysis, which revealed three subpopulations of microglia in the hippocampus (**Figure 7A**). None of the subpopulations showed higher abundance in either genotype. The subpopulations also did not segregate based on expression of established microglial profile markers, such as homeostatic, disease-associated microglia (DAM) (*56*), type I interferon-induced (IFN), and major histocompatibility complex-expressing (MHC) signatures (*42*) (**Figure 7B**). Expression of homeostatic markers was high in all microglia subpopulations and did not change in microglia from the hippocampus of Dutch mice (**Figure 7C**). In addition, no differences were seen in IBA1 IHC between 12-month-old Dutch and wildtype mice (**Supplemental Figure 13A-B**). We observed an approximately 3-fold increase in co-localization of C1q with synaptic puncta in Dutch mice (p = 0.0181, **Supplemental Figure 13C**). These data strengthen the conclusion that C1q–synapse association is significantly elevated in Dutch mice at 12 months of age, as was reported in J20 mice (*57*).

**Figure 7.**
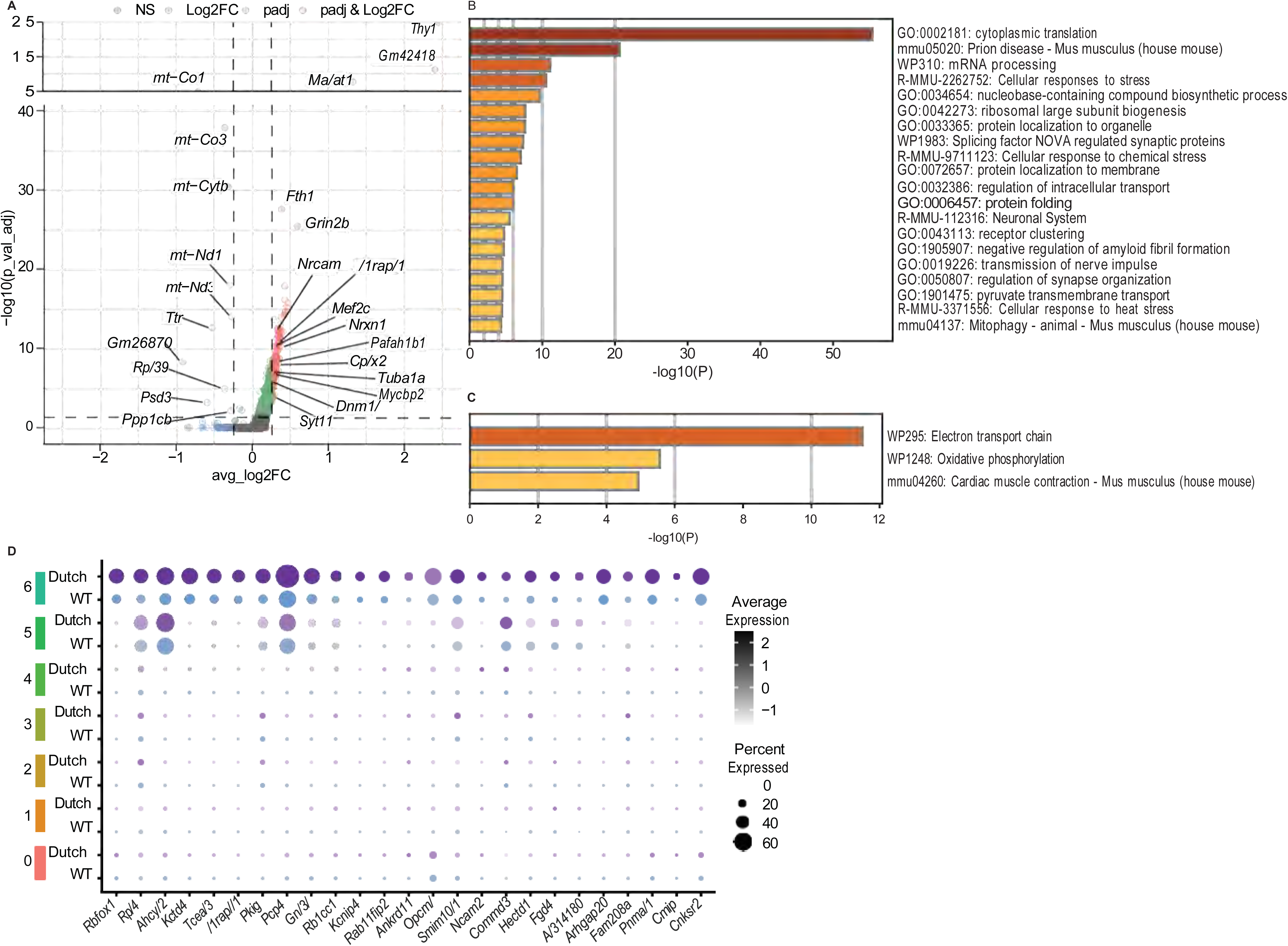
Transcriptional profile of Cluster 6 excitatory neurons is altered in Dutch mice. **(A)** Volcano plot of genes enriched in Cluster 6 as calculated by pseudobulk differential expression analysis. Red genes are Log2FC > 0.25 and adj. p-value < 0.05, blue Log2FC < 0.25 and adj. p-value < 0.05, green genes are Log2FC > 0.25 and adj. p-value > 0.05, and grey genes are Log2FC < 0.25 and adj. p-value > 0.05. **(B-C)** GO term analysis of **(B)** upregulated and **(C)** downregulated genes enriched in Cluster 6 (adj. p-value < 0.05) using metascape(*91*). **(D)** Dot plot of DEGs unique to Cluster 6.

### Dutch mice show changes in gene expression associated with translation in excitatory neurons

Cluster 6 was identified as excitatory neurons and showed the most extensive and unique transcriptional response in Dutch mice (**Figure 6C-D**, **Figure 7A**). In the GO term analysis, upregulated Cluster 6 DEGs included “cytoplasmic translation” (*Eif2s2, Eef1a1, Eef1b2, Rpl22, Rpl26, Rpl27*) and “negative regulation of amyloid fibril formation” (*Cryab, Pfdn6, Slc25a4, Grin2b, Stmn1, Smad4, Pik3r1*, **Figure 7B, Supplemental Table 6**). Notably, “regulation of synapse organization” was also among the top GO terms (*Cask, Cdh8, Gpc4, Grin2b, Mef2c, Nrxn1, Pafah1b1, Ptprd, Tuba1a, Ube2v2, Malat1, Opa1, Mycbp2, Nrcam, Il1rapl1, Arhgef9*), a selection of which are also highlighted in **Figure 7A**. In the GO term analysis, downregulated Cluster 6 DEGs included those related to processes associated with mitochondrial bioenergetic function: i.e., “electron transport chain” (*Mt-Co1, Mt-Co3, Mt-Cytb, Ppp1cb, Calm2*) and “oxidative phosphorylation” (*mt-Atp6, mt-Nd1, mt-Nd3)* (**Figure 7C, Supplemental Table 7**). A selection of unique DEGs from Cluster 6 is shown in **Figure 7D**.

Gene expression was also examined for biological processes associated with presynaptic mechanisms and synaptic transmission, specifically “regulation of synapse organization”, which is a subset of “regulation of cell morphogenesis” (**Supplemental Figure 14A**). Myocyte Enhancer Factor 2C (*Mef2c*) is among this gene list and has been identified as a candidate risk gene for developing late onset Alzheimer’s disease (LOAD) (*58, 59*). Considering that oligodendrocytes can modulate presynaptic properties and synaptic transmission (*60, 61*), relevant biological processes such as “modulation of chemical synaptic transmission” were assessed and found among the top biological processes in Cluster 4 oligodendrocytes (**Supplemental Figure 14B, Supplemental Table 8**). Approximately 25% of the genes present in this biological process are among DEGs in Cluster 6. Genes associated with synaptic transmission that were among those specific to Cluster 4 included transferrin (*Trf,* a recently identified biomarker of AD) (*62*), and carbonic anhydrase 2 (*Car2*). Carbonic anhydrase inhibitors have been considered for prevention of neurovascular dysfunction in CAA and AD due in part to their ability to prevent mitochondrial dysfunction and oxidative stress (*63, 64*).

A cell-cell communication analysis was performed using the software tool Cellphone DB to compute interactions between clusters based on the known Ligand-Receptor (LR) pairs (**Supplemental Table 9**). For each LR in each sample, the software estimated the interaction strength from Cluster 4 oligodendrocytes to Cluster 6 excitatory neurons, from which differences in interaction strengths between Dutch and WT mice were evaluated by t-test. Twelve LR pairs showed significant interaction difference at p-value ≤ 0.05, and all twelve presented higher interactions in Dutch mice. Among these LRs, *Gabbr2* was up-regulated in the Cluster 6 excitatory neurons in Dutch mice compared to those of wildtype (log2FC = 0.19, adj. p-value = 3.6E-4). In Dutch compared to wildtype mice, stronger interactions were exhibited by *Gabbr2* with multiple GABA inhibitory neurotransmitter complex partners including GAD2, SLC6A11, SLC6A1, SLC6A8, SLC6A6, and SLC32A1.

### Dutch mice have fewer and smaller mitochondria in CA1 presynaptic boutons

The number and morphology of mitochondria were assessed in the presynaptic terminals of excitatory neurons in the *stratum radiatum* of Dutch and wildtype mice (**Figure 8A, B**). Previously, NFA-Aβ has been reported to cause imbalance in mitochondrial fission-fusion, leading to fragmented morphology and altered function (*65–69*). Fewer presynaptic mitochondria were found in Dutch compared to wildtype mice (**Figure 8C**, mean ± SD for total of mitochondria per presynaptic bouton = 75.60 ± 14.62 in Dutch and 106.3 ± 20.06 in wildtype, p = 0.0324 by two-tailed unpaired t test). This deficiency in the Dutch mice was likely due to fewer straight mitochondria at the presynaptic terminals, because curved mitochondria numbers were unchanged between genotypes (**Figure 8D-F**, mean ± SD for number of straight mitochondria = 71.80 ± 15.06 in Dutch and 101.3 ± 17.95 in wildtype, p = 0.314 by two-tailed unpaired t test. Mean ± SD for number of curved mitochondria = 3.80 ± 2.59 in Dutch and 5.00 ± 2.16 in wildtype). In assessments of differences in morphology, mitochondrial diameters were comparable (**Figure 8G**, mean ± SD = 301.8 ± 28.73 in Dutch and 328.6 ± 19.29 in wildtype), but shorter lengths were found in 12-month-old Dutch compared to wildtype mice, as estimated from the number of layers traversed in the consecutive sections by each mitochondrion (**Figure 8H**). Each layer traversed was ∼ 80 nm using 3-dimensional reconstruction of measurements from serial sections.

**Figure 8.**
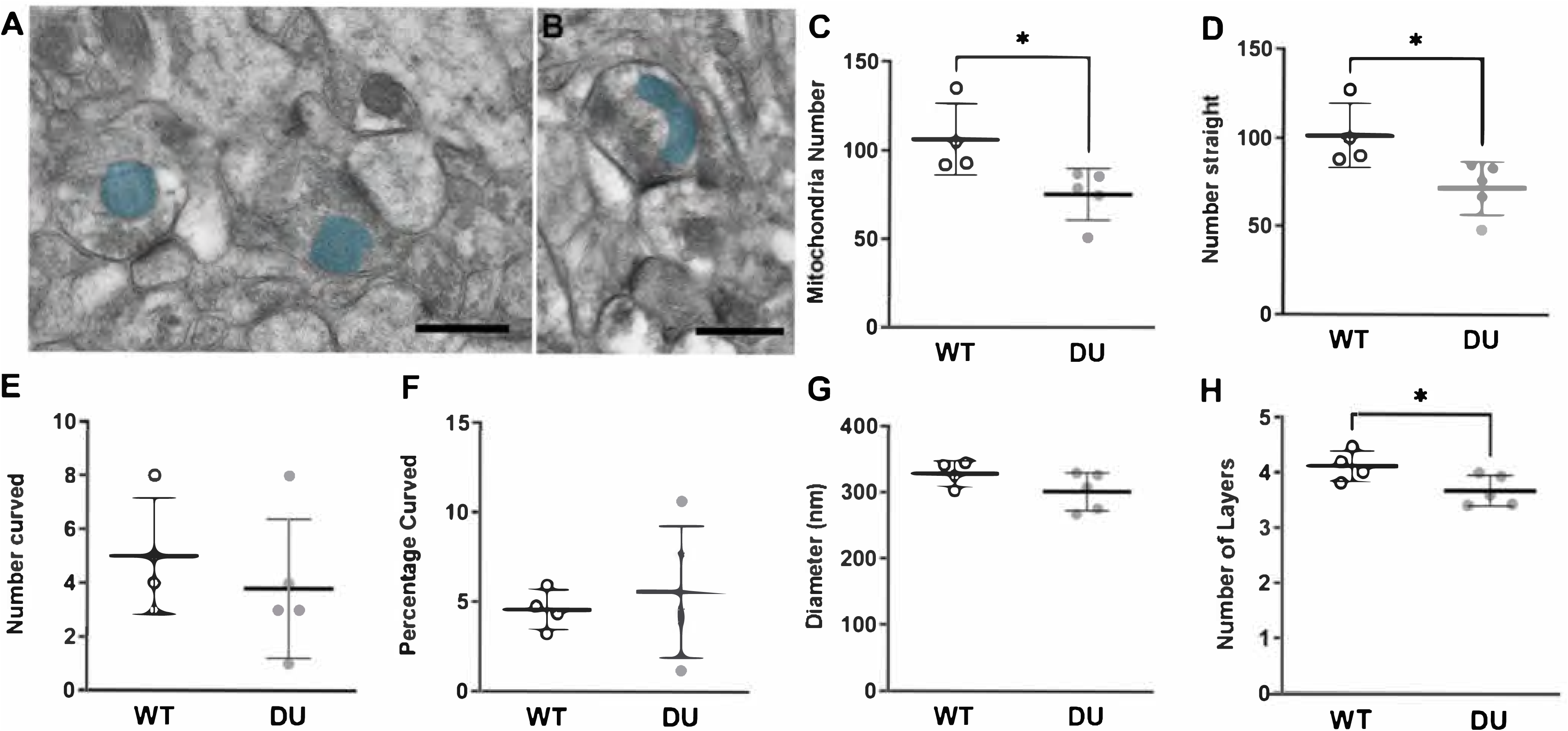
Fewer mitochondria were present in presynaptic boutons at excitatory synapses in the CA1 of 12-month-old Dutch mice compared to wildtype. **(A)** Electron micrographs showing representative examples of presynaptic mitochondria (blue masks). **(B)** Curved presynaptic mitochondria demonstrate a bend of at least 90°. Straight presynaptic mitochondria did not exhibit any curvature. Scale bar = 500 nm. **(C)** Total number, **(D)** number of straight mitochondria, **(E)** number of curved mitochondria, **(F)** percentage of curved mitochondria, **(G)** average diameter, and **(H)** number of layers traversed by each mitochondrion in wildtype and Dutch mice. Data are mean ± SD from n = 4 wildtype and n = 5 Dutch mice.

### Dutch mice have reduced mitochondrial electron transport chain complex I activity

The activities of electron transport chain complexes I-IV and citrate synthase were assessed in hippocampi from 7- and 12-month-old wildtype and Dutch mice as described (*70, 71*). Each complex activity was standardized to either μg protein (p = 0.0355) or to mU/ml of citrate synthase (CS) and analyzed using an unpaired t test (p = 0.0197) (**Supplemental Figure S15, Supplemental Tables 10, 11**). Mitochondrial complex I activity was reduced in 12- but not 7-month-old Dutch compared to that of wildtype mice.

### Dutch and Arctic Aβ peptides increase mitochondrial membrane potential in cultured cells

Both Dutch APP^E693Q^ and Arctic APP^E693G^ are known to form primarily oligomers. We assayed wildtype, Dutch, and Arctic mutant Aβ peptides (**Supplemental Figure S16**) using cultured Neuro2a cells. In this experiment, Neuro2a cells were respectively treated for 24 h with wildtype peptide and Aβ carrying at position 22 either the Dutch E693Q or the Arctic E693G mutation. Cells were then stained with a MT-1 MitoMP detection kit, using a mitochondrial membrane potential-sensitive dye, and MitoTracker Green, which labels total mitochondrial mass. Representative images are presented from one of three independent experiments showing similar results. For quantitative analysis, 30 cells per condition were analyzed. Mitochondrial membrane potential was increased by both oligomerogenic mutant peptides, as reported (*72*). Data were analyzed by one-way ANOVA followed by Dunnett’s post hoc test. NS, not significant (p > 0.05); ****, p< 0.001. Scale bar, 10 μm.

## Discussion

The localization of mouse brain Dutch NFA-Aβ was ascertained using immunohistochemistry with A11 antibody and fluorescence microscopy with a cyclic peptide-FITC ligand (*22*). In contrast to the apparently normal density and morphology of synapses in excitatory neurons, we observed abnormalities in the apical dendrite structure in the CA1 of Dutch mice (*24*). Impaired short-term synaptic plasticity developed at the CA1-CA3 synapses first appearing at the age of 11 to 13 months.

Assessment of the I-O curve and the LTP response after high frequency stimulation in the CA1-CA3 synapses, indicated no difference between Dutch and wildtype genotypes. In contrast, intracerebroventricular injection of synthetic Dutch and Arctic (APP^E693G^) Aβ peptides have been previously reported to block LTP in rats (*73*). Despite normal *long-term* synaptic plasticity in young or old Dutch mice, defects in several types of *short-term* plasticity emerged in the aging 11- to 13-month-old Dutch mice. Specifically, mice carrying the Dutch mutation exhibited impaired PTP, synaptic fatigue (SF), and impaired replenishment of the synaptic pool, but unaffected PPF. Both PPF and PTP are related to accumulation of Ca^2+^ at the presynaptic terminal, due (respectively) to two closely paired pulses or to high frequency stimulation, both of which augment neurotransmitter release. PPF and PTP are governed by different synaptic vesicle proteins: e.g., the absence of the synaptic vesicle protein synaptogyrin I in mice causes respectively impaired PTP but no effect on PPF (*74*).

Dutch mice showed altered presynaptic plasticity mechanisms related to the RRP of neurotransmitters. Dutch mice exhibited respectively faster rates of decay and recovery during high frequency stimulation that causes fatigue at the synapses and after synaptic depression. The depletion and replenishment of the RRP of neurotransmitters appear more profound in Dutch mice. The RRP of neurotransmitters refers to a small subset of neurotransmitter-containing vesicles residing close to the presynaptic terminal (either docked or non-docked) and disposed for immediate released during stimulation. Several parameters can affect the decay of the RRP; e.g., replenishment of the RRP could be mediated by the Ca^2+^-mediated recruitment of vesicles from the reserve pool to the RRP.

The required interplay between pre- and postsynaptic neurons during synaptic plasticity suggests that presynaptic changes could affect activation of postsynaptic neurons eventually leading to long-term synaptic impairments. High frequency stimulation resulted in normal LTP, suggesting that Dutch mice do not display derangement of postsynaptic mechanisms at the ages studied, an observation that could be attributed to compensatory mechanisms. In one possibility, the decrease in release probability with PTP may be balanced by the increased depletion and replenishment of the RRP.

Hippocampal microglia (Cluster 2) did not show a high number of DEGs, nor were differences in expression of microglia DAM and homeostatic signatures observed between Dutch and wildtype mice. The highest total and unique number of DEGs were found in excitatory neurons (Cluster 6) and were associated with processes such as “protein translation” and “electron transport chain”. LR pairs were examined between oligodendrocytes and excitatory neurons, which indicated that *Gabbr2* showed stronger interaction with multiple GABA inhibitory neurotransmitter complex partners in Dutch mice.

Finally, prior studies using bulk RNAseq on the entorhinal cortex and dentate gyrus of wildtype and Dutch mice were re-examined using scRNA-seq, revealing no obvious differences in microglial transcriptional profiles. Moreover, the Dutch mouse hippocampal transcriptome did not present neuroinflammatory and microglial transcriptomic responses which are commonly observed in models with senile plaques and amyloid fibrils (e.g., PSAPP, 5XFAD). Failure to capture such a response may be due to the cell populations selected for profiling. Indeed, previous characterization indicated that microgliosis in other Dutch mice (*75*) was not generalized but was limited to the immediate proximity of Aβ-laden vessels. In this study, no differences were observed in microgliosis, which is reported to be an NFA-Aβ-induced process that drives activation of complement and synapse elimination (*57*). In a transcriptomic analysis on postmortem brain tissue from humans with HCHWAD, inflammatory genes and pathways were also not prominently represented (*76*). As detected in the Dutch mouse, the main pathways over-represented in the human HCHWAD brains were related to mitochondrial dysfunction (*76*). The principal up-regulated pathways entail interactions of the extracellular matrix and transforming growth factor beta signaling. In a clustering analysis to identify molecular subtypes of AD, gene expression profiles have been gleaned from the Mount Sinai Brain Bank (MSBB-AD) and Religious Orders Study-Memory and Aging Project (ROSMAP) (*77*). The Dutch mouse gene signature significantly correlated with the subtype of “typical” or “Aβ predominant” AD presentation, and was characterized by increased immune activation and inflammation, as well as decreased synaptic signaling and dysregulation in neuronal processes and synaptic signaling (*77*). Taken together, the results minimize the contribution of neuroinflammation in the pathogenesis leading to learning disorder in Dutch mice.

Among the most dysregulated pathways in the Dutch mouse brain was “structural constituent of ribosome”, which is associated with the most significant GO term in Cluster 6 “protein translation”. Increasing evidence suggests that dysregulation of mRNA translation contributes to the pathogenesis of several neurological diseases, including AD (*78*). Accumulation of Aβ is linked to increased protein synthesis through loss of interactions from key translation regulators, particularly translation repressors (*79*). When the cell-type specific transcriptional responses to NFA-Aβ were examined, altered transcriptional profiles were identified in multiple cell types. In these profiles, overlap with GO term enrichment analysis indicated “amyloid-beta binding” among the shared processes.

The Cluster 6 neuron DEGs downregulated in Dutch compared to wildtype mice were associated with mitochondrial functions, specifically GO terms “electron transport chain” and “oxidative phosphorylation”. In concordance, cellular respiration pathways were the most significantly altered pathways identified in the transcriptomic analysis from human patients with HCHWAD (*76*). Moreover, fewer and smaller mitochondria were found at the presynaptic terminals of excitatory neurons in the CA1 of Dutch compared to wildtype mice. Strong evidence linking NFA-Aβ to mitochondrial complex I endpoints has come from the combined use of a genetically encoded fluorescent redox sensor and multiphoton microscopy to image and monitor *in vivo* the mitochondrial redox imbalance in neurons from the APP/PS1 transgenic mouse model of AD (*80*). The deposition of Aβ plaque and particularly soluble NFA-Aβ species were shown to lead to increased mitochondrial oxidative stress levels within neuronal mitochondria (*80*). Mitochondrial redox dysregulation in neurons was prevented by inhibition of mitochondrial calcium influx, which is pathologically exacerbated in Aβ plaque-depositing mice (*80*). Furthermore, mitochondria-targeting antioxidants decreased mitochondrial oxidative stress back to basal levels and reverted Aβ plaque-associated dystrophic neurites in the APP/PS1 mice (*80*). Defective mitochondrial complex I activity has been detected in living human AD brain (*81*). Altered mitochondrial fission and fusion has been observed in the hippocampus of human AD and in Tg2576 mice and may explain the observations of decreased mitochondrial density and smaller mitochondria in Dutch mice (*68, 82*). Increased mitochondrial fragmentation has been reported in neurons exposed to NFA-Aβ (*83*). Elevated levels of NFA-Aβ impaired transport of mitochondria in dendrites and axons and increased mitochondrial permeability in mice harboring the Osaka mutation of APP (APP^E693D^) (*84, 85*). Mitochondria-targeting therapy holds promise for treating neurodegenerative disease (*86, 87*). Our finding of reduced density of presynaptic mitochondria in the CA1 of Dutch mice suggests that NFA-Aβ might also affect transport of mitochondria to axon terminals. The dysfunction of mitochondria and endoplasmic reticulum (ER) protein translation observed in Dutch mice is consistent with observations in other mouse models of AD (*88*).

In conclusion, two independent methods visualized for the first time *endogenously generated* NFA-Aβ *in vivo* in the absence of intracerebral injection of synthetic peptides and without the concurrent presence of Aβ fibrils. Changes related to the *APP^E693Q^*-genotype were revealed in NFA-Aβ histopathology, in expression of mitochondrial and neuronal translation-related genes, in presynaptic physiology, and in the ultrastructure of presynaptic mitochondria. Neuroinflammation is not a major contributor to the pathogenesis in Dutch mice. Neither fibrillar Aβ nor neuroinflammation is required for the aging-related learning behavior phenotype.

This study of APP^E693Q^ mice is particularly timely considering the promising outcome of clinical trials for lecanemab (*89, 90*) and donanemab (*91*). After 18 months of treatment with antibody, participants experiencing mild cognitive impairment due to AD showed respectively a modest but statistically significant 27% and 35% slower rate of cognitive decline with lecanemab and donanemab. The modest clinical response to therapeutic infusion of anti-amyloid antibodies may be attributable, at least in part, to the persistence of residual toxic NFA-Aβ that goes undetected by current fluid biomarker analysis and amyloid fibril imaging. The potential clinical importance of NFA-Aβ is reinforced by the recent discovery of a new FAD mutation (*APP^D678N^*) that also favors NFA-Aβ accumulation (*92*). Therapeutic depletion of NFA-Aβ may be required for optimum and more robust cognitive benefits from anti-amyloid antibody infusion. Newer methods for imaging NFA-Aβ (e.g., [^64^Cu]-NOTA cyclic D,L-α-peptides) in the living human brain (*22*) and assays for detection of NFA-Aβ in biofluids (*93–95*) may provide earlier diagnosis and improved monitoring of patients undergoing Aβ immunodepletion therapy by assuring that both fibrillar and nonfibrillar Aβ conformers are cleared as completely as possible.

## Supporting information

Supps

## Acknowledgments

We would like to thank all the members of the Dickstein, Arancio, Levy, Rahimipour, Lubell, Guerin, and Ehrlich/Gandy laboratories, and William G. M. Janssen, Allison Sowa, Daniel R. Dickstein, Frank Yuk, and Michael Ragusa for technical assistance and discussion. The authors would like to thank Yanzhuang Wang (Shenzhen Bay Laboratory), and Gary Gibson, Giovanni Manfredi, and Csaba Konrad (all from Weill Cornell Medical College) for valuable conversations and suggestions. We also thank Tamas Kozicz and Graeme Preston (Icahn School of Medicine at Mount Sinai) for performing the electron transport chain activity assays.

## Conflicts

Dr. Gandy is a co-founder of Recuerdo Pharmaceuticals. He has served as a consultant in the past for J&J, Diagenic, and Pfizer, and he currently consults for Cognito Therapeutics, GLG Group, SVB Securities, Guidepoint, Third Bridge, MEDACORP, Altpep, Vigil Neurosciences, Memory Garden, and Eisai. He has received research support in the past from Warner-Lambert, Pfizer, Baxter, and Avid. The authors declare no other competing interests.

## Funding

National Institutes of Health grant F32AG077900 (ELC)

National Institutes of Health grant P30 AG066514 to Mary Sano with Developmental Pilot Award (ELC, SG)

National Institutes of Health grant U01AG046170 (SG, MEE, MW, BZ)

National Institutes of Health grant RF1AG058469 (SG, MEE)

National Institutes of Health grant RF1AG059319 (SG, MEE)

National Institutes of Health grant R01AG061894 (SG, MEE)

Cure Alzheimer’s Fund (SG, MEE, SR, BG, WL)

National Institutes of Health grant RF1AG057440 (MW, BZ)

National Institutes of Health grant RF1AG056507 (CG)

National Institutes of Health grant AG056732 (EL)

National Institutes of Health grant RF1AG086510 (EL)

National Institutes of Health grant R01NS110024 (OA)

Alzheimer’s Association NIRG-12-242386 (DLD)

Medical Student Training in Aging Research Program Scientific Fellowship from the American Federation for Aging Research (EB)

The I. M. Ertegun Project for Cognitive Health Extension (SG, MEE)

National Institutes of Health grant RF1AG077828 (MW)

National Institutes of Health grant R21AG077168 (MW)

Alzheimer’s Association AARG-22-928419 (MW)

AMED under Grant Number JP24wm0625509 (KA)

Grant-in-Aid for Scientific Research (B) [24K02860] (KA)

**Consent** was unnecessary. No human subjects were involved in this study.

## Author contributions

Conceptualization: MEE, SG, EL, DLD, OA

Methodology: EB, MV, DLD, KT, SH, JVDL, OA, EA, ELC, CDS, MW, BZ, CG, SR, YH

Investigation: RT, MV, DL, EB, DLD, FG, ELC, KT, SH, JVDL, OA, EA, HZ, BZ, MW, CDS, LS, LL, MI, SR, AA, KA, TS, YH

Visualization: MV, DLD, KT, SH, ELC, MW

Supervision: MEE, SG, EL, DLD, OA

Writing—original draft: MV, DLD, EB, ELC, SG, EA, OA, MW

Writing—review & editing: MV, DLD, ELC, SG, MEE, EL, CG, SR, WL, BG, TS, YH

## Data and materials availability

The scRNA-seq data (cell ranger processed count matrices) are in a Synapse repository (accession syn52752404, with direct link https://www.synapse.org/#!Synapse:syn52752404/files/). All data are available in the main text or the supplementary materials.” The APP^E693Q^ transgenic mouse can be provided by Thomas Jefferson University pending scientific review and a completed material transfer agreement. Requests for the APP^E693Q^ transgenic mouse should be submitted to.

## Disclaimer

The opinions expressed herein are those of the authors and are not necessarily representative of those of the government of the United States, the Uniformed Services University of the Health Sciences, the Department of Defense (DoD), or the United States Army, Navy or Air Force or the Henry M. Jackson Foundation for the Advancement of Military Medicine, Inc.

## Supplementary Materials Table of Contents

Supplementary Methods

Figs. S1 to S16

References (92 to 102)

## Supplementary Tables Titles and Captions

**Supplementary Table 1. Antibodies used in this study.**

**Supplementary Table 2. Oligomer level and electrophysiological data analyzed by sex.**

**Supplementary Table 3. Cluster Cell Numbers.** Number of cells per cluster per sample.

**Supplementary Table 4. Cluster Markers.** De novo cluster signatures calculated by comparing cell clusters using Wilcox rank sum test in Seurat.

**Supplementary Table 5. Cluster DEGs.** Differentially expressed genes across all 14 clusters using MAST implemented in Seurat.

**Supplementary Table 6. Cluster 6 GO-terms.** GO-term analysis of genes enriched in Cluster 6 (l2f>0, padj <0.05) using metascape.

**Supplementary Table 7. Cluster 6 GO-terms.** GO-term analysis of genes enriched in Cluster 6 (l2f<0, padj <0.05) using metascape.

**Supplementary Table 8. Cluster 4 GO-terms.** GO-term analysis of genes enriched in Cluster 6 (l2f>0, padj <0.05) using metascape.

**Supplementary Table 9. Ligand-Receptor Pairs for Clusters 6 and 4.** Cell-cell communication analysis calculated by Cellphone DB to compute interaction between Clusters 6 and Clusters 9 based on known Ligand-Receptor pairs. t-test was used to compare interaction strength differences between Dutch and WT mice

**Supplementary Table 10. Mitochondrial electron transport chain (mETC) enzymology. mETC complexes standardized to µg protein.**

**Supplementary Table 11. Mitochondrial electron transport chain (mETC) enzymology. mETC complexes standardized to mU/ml citrate synthase.**

## Highlights

- Transgenic *APP^E693Q^* mice develop learning behavior changes associated with accumulation of nonfibrillar aggregates of Dutch Aβ in their brains.
- Electrophysiology shows aging-related defects in post-tetanic potentiation, synaptic fatigue, and synaptic vesicle replenishment in *APP^E693Q^* mice.
- Single cell transcriptomics did not identify the signature of disease-associated microglia, but the significant association of C1q with synaptic puncta indicates activation of the innate immunity pathway leading to microglial engulfment of synapses.
- Ultrastructural analysis showed that Dutch mice have fewer and smaller mitochondria in CA1 presynaptic boutons. Excitatory neurons show changes in endoplasmic reticulum- and mitochondria-related transcripts. Activity of mitochondrial complex I (but not complexes II, III, or IV) is diminished in an aging-related fashion in the *APP^E693Q^* mouse hippocampus.

## RESEARCH IN CONTEXT

1. **Systematic review**: The authors reviewed previous publications using traditional sources, including PubMed and preprint server databases. Nonfibrillar aggregates (NFAs), oligomers, and protofibrils have been frequently studied using synthetic peptides or cell media. There are relatively few peer-reviewed publications regarding endogenously generated nonfibrillar aggregates of Aβ in the brain and/or within the context of Alzheimer’s disease (AD). We have cited relevant publications.
2. **Interpretation**: Our results show that aging-related accumulation of NFA-Aβ in *APP^E693Q^* mice could be visualized using A11 immunohistochemistry or NFA-Aβ-detecting cyclic D,L-α-peptide-FITC microscopy. The presynaptic termini of *APP^E693Q^* mice developed aging-related physiological abnormalities in post-tetanic potentiation, synaptic fatigue, and synaptic vesicle replenishment. Single-cell RNA sequencing showed that excitatory neurons exhibited the most altered transcriptomic profile, especially involving “protein translation” and “oxidative phosphorylation”. Direct measurements of electron transport chain catalysis revealed an aging-related reduction in mitochondrial complex I activity in the hippocampi of Dutch mice. Microglial transcript analysis revealed no evidence of inflammation.
3. **Future directions**: Accumulation of endogenously generated Dutch NFA-Aβ can be visualized with new ligands and alters neuronal metabolism but does not activate inflammation. Since endogenously generated NFA-Aβ is associated with learning behavior changes, the depletion or neutralization of <u>*both fibrillar and NFA-Aβ*</u> may be needed for complete elimination of Aβ toxicity when anti-amyloid antibodies are used clinically.

## Notes

https://www.synapse.org/#!Synapse:syn52752404/files/

### Summary of Updates

Updated with additional experiments, figures, and data

