## Supplementary material for "Endogenously generated Dutch-type Aβ nonfibrillar aggregates dysregulate presynaptic neurotransmission in the absence of detectable inflammation": Supps

Click here to access/download
**Supplementary files**
CASTRANIO SUPP TABLE 1_020926_SG_word version.docx

Click here to access/download

**Supplementary files**

Supplemental Tables 2-6_020926_SG.xlsx

Click here to access/download
**Supplementary files**
Supp Table 7.pdf

Click here to access/download
**Supplementary files**
Castranio_Supplemental Table 8_012025_SG.docx

Click here to access/download
**Supplementary files**
Castranio_Supplementary_Table 9_012025_SG.docx

Click here to access/download
**Supplementary files**
CASTRANIO SUPP TABLE 10_020326_SG_word version.docx

Click here to access/download
**Supplementary files**
CASTRANIO SUPP TABLE 11..pdf

Supplementary Figure 1

Click here to access/download
**Supplementary files**
Castranio_Supp_Figure_1_032526_SG_C.pdf

Supplementary Figure 2

Click here to access/download
**Supplementary files**
Castranio_Supp_Figure_2_032526_SG_C.pdf

Supplementary Figure 3

Click here to access/download
**Supplementary files**
Castranio_Supp_Figure 3_032526_SG_C.pdf

Supplementary Figure 4

Click here to access/download
**Supplementary files**
Castranio_Supp_Figure 4_032526_SG_C.pdf

Supplementary Figure 5

Click here to access/download
**Supplementary files**
Castranio_Supp_Figure 5_032526_SG_-C.pdf

Supplementary Figure 6

Click here to access/download

**Supplementary files**

Castranio_Supp_Figure 6_032526_SG_-C.pdf

Supplementary Figure 7

Click here to access/download
**Supplementary files**
Castranio_Supp_Figure 7A-I_032526_SG_C.pdf

Supplementary Figure 8

Click here to access/download
**Supplementary files**
Castranio_Supp_Figure 8_0325226_SG_C.pdf

Supplementary FIgure 9

Click here to access/download
**Supplementary files**
Castranio_Supp_Figure 9_032526_SG_C.pdf

Supplementary Figure 10

Click here to access/download
**Supplementary files**
Castranio_Supp_Figure 10_032526_SG-C.pdf

Supplementary Figure 11

Click here to access/download
**Supplementary files**
Castranio Supp Fig 11 032526_SG_Clean.pdf

Supplementary FIgure 12

Click here to access/download
**Supplementary files**
Castranio_Supp_Figure 12_032526_SG_-C.pdf

Supplementary Figure 13

Click here to access/download
**Supplementary files**
Castranio_Supp_Figure 13_Clean_022226_SG.pdf

Supplementary Figure 14

Click here to access/download

**Supplementary files**

Castranio_Supp_Figure 14_032526_SG_C.docx

Supplementary Figure 15

Click here to access/download
**Supplementary files**
Castranio_Supp_Figure 15_032526_SG_C.docx

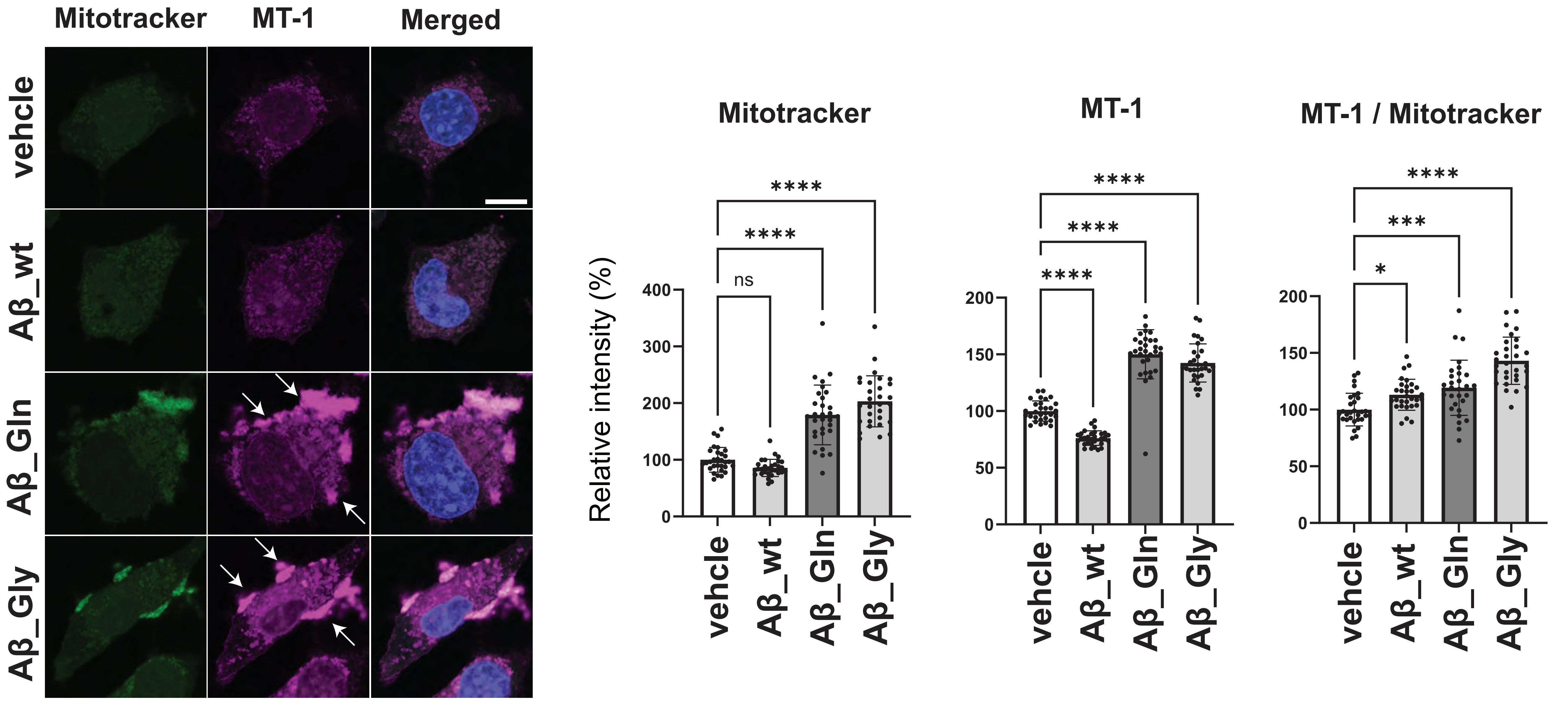

Mitotracker
MT-1
Merged
vehcle
Aβ_wt
Aβ_Gln
Aβ_Gly
Mitotracker
MT-1
MT-1 / Mitotracker
Relative intensity (%)
ns
****
***
*

Click here to access/download
**Supplementary files**
Castranio Supp Meth Results
032926_SG_Clean.docx.pdf
